# Antigen presenting cells play a critical role in determining response to regulatory T cell therapy in type 1 diabetes

**DOI:** 10.1101/2025.11.15.688620

**Authors:** Mackenzie Dalton, Emmanuel Asante-Asamani, James M. Greene

## Abstract

Type 1 diabetes (T1D) is an autoimmune disease in which the immune system attacks pancreatic beta cells, leading to high blood glucose levels and requiring lifelong insulin therapy. There is no cure, and individuals with T1D may face a reduced lifespan of up to 12 years. Defects in regulatory T cells (Tregs) are a key contributor to disease onset and are being explored as a therapeutic avenue. However, the effectiveness of Treg therapy remains uncertain. Research is further limited by the inability to directly observe pancreatic and lymph node activity during the long presymptomatic stage of T1D. In this study, we develop a mathematical model for beta and T cell dynamics. We find both Treg quality and quantity affect disease progression, and that antigen-presenting cell (APC) dynamics play a central role. Notably, Treg therapy combined with APC depletion improves outcomes, especially with strong peptide-induced APC activation.

## 1 Introduction

Type 1 diabetes (T1D) is an autoimmune disease in which the immune system attacks the insulin-producing pancreatic beta cells, leading to elevated glucose levels and the inability of the body to regulate blood sugar naturally. Although exact causes are unknown, both genetic and environmental factors are believed to contribute to its onset, with the concordance rate between monozygotic twins reported to be somewhere between 50-70% [1, 2]. Furthermore, it is one of the most common autoimmune disease in children [3]. Currently, there is no cure for T1D, and patients with the disease require lifelong daily insulin injections and glucose monitoring. However, despite the availability of treatment, patients still face heightened risks of complications, including heart disease, kidney failure, and stroke [4]. Moreover, patients with type 1 diabetes face an average reduction in lifespan of 12 years compared with their healthy counterparts, emphasizing the need for a cure [5].

The progression of type 1 diabetes occurs in three stages. Initially, a patient tests positive for two or more autoantibodies associated with type 1 diabetes, yet maintains normal glycemic levels and remains asymptomatic. In this initial stage, most patients are unaware that they are developing type 1 diabetes unless they are tested for the presence of autoantibodies [6]. The four types of autoantibodies that are typically found in patients with T1D are islet cell cytoplasmic autoantibodies (ICA), glutamic acid decarboxylase autoantibodies (GADA), insulin autoantibodies (IAA), and insulinoma-associated-2 autoantibodies (IA-2A) [7]. Studies have shown that testing positive for two or more autoantibodies associated with type 1 diabetes leads to an almost guaranteed diagnosis of type 1 diabetes within 10 years [8]. Moreover, by 2 years of age, the presence of autoantibodies is highly predictive of disease. In fact, it is estimated that 90% of children who develop disease before the age of puberty have detectable autoanitbodies before the age of 5 [9]. The second stage of T1D progression is defined as when a patient has two or more autoantibodies together with abnormal glucose levels due to a decrease in beta cell mass; however, in most cases the individual remains asymptomatic at this stage of disease. The final stage of T1D progression is when clinical diagnosis occurs and T1D symptoms are present. During this stage, substantial beta cell mass has been lost, and insulin therapy is required for survival [10].

Regulatory T cells (Tregs) play a crucial role in maintaining immune tolerance and preventing an autoimmune response [11, 12]. For example, in IPEX (immune dysregulation, polyendocrinopathy, enteropathy, X-linked) syndrome, regulatory T cells are either missing or dysfunctional, which leads to numerous health problems, including the early onset of type 1 diabetes, with more than 80% of patients developing T1D before the age of 2 [13, 14]. While Tregs are known to play a role in T1D, it remains to be understood what role *quantity* versus *quality* of regulatory T cells plays in disease progression. More precisely, in T1D the quantity (i.e. number) of regulatory T cells has been reported to be unaltered in some studies [12, 15] and reduced in others [15]. Conversely, the quality of Tregs refers to their suppressive capabilities. The immune system is naturally equipped with mechanisms to manage hyperactivity and self-regulation; regulatory T cells act as mechanism of “immunological tolerance” [16], and can prevent autoimmunity through several pathways [17]. However, while there are a plethora of functions of regulatory T cells, their precise mechanism of action in vivo remains poorly understood [18]. In this work, we investigate two potential methods in which regulatory T cells suppress effector T cell activation and capabilities. First, Tregs suppress the activation of effector T cells [19]; this occurs, for example, via interactions with dendritic cells in a CTLA4 - dependent manner [20]. Secondly, regulatory T cells also suppress the killing capabilities of effector T cells through the impairment of granule exocytosis, a major mechanism used by effector T cells to induce apoptosis of target cells [14, 21]. We note that it is well-known that patients with T1D exhibit decreased functional capacity in their Treg population [15], and that the quality-versus-quantity question of regulatory T cells is relevant to recent clinical trials in T1D, where Treg therapy failed to demonstrate improved beta cell function [22].

Mathematical modeling is a crucial tool for exploring immune dynamics during the prolonged presymptomatic phase of type 1 diabetes, where most beta cells are destroyed prior to clinical diagnosis. This extended period complicates early detection and limits direct studies of disease progression, which often rely on cadaveric analysis due to the pancreas’s challenging location. By providing a framework to investigate T1D progression, mathematical models help elucidate the role of regulatory T cells in T1D. There are many mathematical models that study the progression of type 1 diabetes [23–34]; however, few of these models incorporate regulatory T cells and their mechanisms of action [23,24,29,33,34]. We next discuss previous mathematical models that incorporate Treg dynamics, and highlight aspects that have not been considered regarding the quality-versus-quantity question.

In [34], the authors are primarily interested in the role of viral infections in triggering T1D, which is not a focus of this work. While the model developed by Nelson et al. [23] is able to study both regulatory and effector T cells during disease progression, it does not include antigen-presenting cells (APCs), an essential component of immunological tolerance that activates both Tregs and effector T cells [35]. In 2019, Shtylla et al. [24] introduced a model that incorporates both regulatory T cells and APCs; however the authors restrict dynamics to the pancreas (i.e. it is a single compartment model). Because of this restriction, the different mechanisms utilized by regulatory T cells are unable to be properly investigated, which was noted by the authors as being a limitation of their study [24]. Indeed many models overlook the significance of distinguishing between the pancreas and lymph nodes as separate modeling compartments, despite the unique regulatory T cell dynamics within each [36]. While the two compartments introduce complexity, this differentiation is vital for fully capturing immune interactions and advancing our understanding of T1D. In particular, studying the different regulatory T cell mechanisms in both the lymph nodes and pancreas are important for understanding and predicting responses to regulatory T cell therapy, a safe and promising therapeutic option for type 1 diabetes patients [37]. We note that the authors of [29] consider similar components as the model we propose below, but utilize an agent-based framework, making it less analytically tractable when compared to the work presented here. Similarly, the authors in [33] construct a high-dimensional ODE system, which prevents any analytical results from being obtained.

In this work, we propose a two compartment mathematical model describing immune dynamics in both the lymph node and pancreatic compartments. Distinguishing between these two compartments allows us to study the two distinct mechanisms of regulatory T cell action. Since activation of effector T cells (as well as Tregs) by APCs occurs primarily in the lymph nodes, we utilize this compartment to study the mechanism of Treg suppression on effector T cell activation by APC [38]. As the the apoptotic function of effector T cells occurs in the pancreas (where the beta cells are located), we investigate the suppression of effector T cell induced apoptosis by regulatory T cells here [39]. Thus our model allows us to distinguish between two of the main mechanisms implemented by regulatory T cells to suppress the immune system, and can thus offer insights into the role of each mechanism. Moreover, by simulating various patient-specific parameter regimes, we utilize the model to elucidate the conditions under which Tregs fail to prevent autoimmune diabetes and explore potential therapeutic strategies to enhance regulatory T cell efficacy. Furthermore, the model also provides insights into why, in some cases, regulatory T cell therapy fails to prevent disease onset. Key biological processes, including regulatory T cell activation, suppression of autoreactive T cells, and feedback mechanisms within the immune system are incorporated. Our findings could be used to provide insights into the optimal modulation of regulatory T cell activity to prevent and/or treat type 1 diabetes, including the possible combination of Treg therapy with APC depletion therapy, which we explore with our proposed model.

## 2 Results

The primary model considered consists of seven ordinary differential equations (ODEs), which are given below:

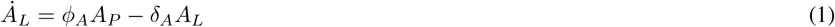

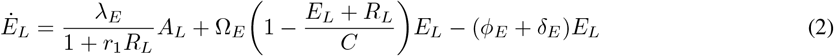

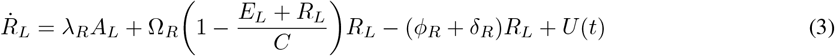

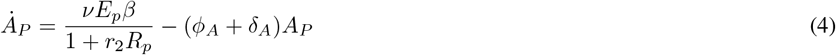

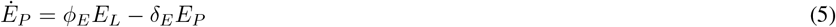

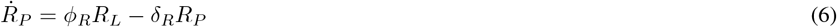

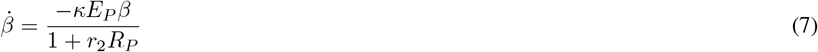

All parameters in the above system are positive, and we term this model the *type 1 diabetes immune* (T1DI) model. We note that equations (1) - (7) are obtained as a quasi-steady state approximation of a model which also incorporates peptide dynamics; see Section 4.1 for details. Below we describe the biological assumptions underlying the above T1DI model.

In (1) - (7), a state variable subscript *L* indicates a population restricted to the lymph nodes, while *P* corresponds to a population in the pancreas. The parameters *ϕ*_*A*_, *ϕ*_*E*_, and *ϕ*_*R*_ represent the respective transition rates of activated APCs, effector T cells, and Tregs between the pancreas and lymph node compartments. Upon activation from beta cell death (*νE*_*P*_ *β*) in the pancreas, APCs in the pancreas (*A*_*P*_) are able to travel to the lymph nodes where they function in two ways. First, activated APCs in the lymph nodes (*A*_*L*_) stimulate self-reactive effector T cells (*E*_*L*_). This activation is suppressed by active regulatory T cells (*R*_*L*_). APCs in the lymph nodes (*A*_*L*_) also activate regulatory T cells. Both regulatory and effective T cells are able to proliferate in the lymph nodes before traveling to the pancreas; this proliferation is modeled as logistic growth with a joint carrying capacity *C*. In the pancreas, effector T cells (*E*_*P*_) kill beta (*β*) cells, and regulatory T cells (*R*_*P*_) suppress this killing. Populations are measured in cell counts, and time units are days. The rate *U* represents an external dosing of Tregs into the lymph node, and is used to model Treg therapy; see equation (34) in Section 4.1.3. For more information regarding parameters, see Table 2 in Section 4.2.

The two mechanisms of regulation of the immune response by the regulatory T cells are modeled by parameters *r*_1_ ≥ 0 in (2) and *r*_2_ ≥ 0 in (4) and (7). Specifically, *r*_1_ denotes the degree to which regulatory T cells suppress effector T cell activation by APCs in the lymph nodes, while *r*_2_ quantifies the suppression of apoptotic ability of effector T cells in the pancreas by Tregs. Phenomenologically, both terms represent a per-cell reduction in the corresponding rate.

At the start of all simulations (*t* = 0 days), stage 1 of the disease is assumed, i.e. there exist overt immunological abnormalities but normal insulin release. Specifically, we assume an initial event, such as a viral infection, led to non-zero initial conditions for both *E*_*L*_ and *R*_*L*_; note that we are not modeling disease initiation in this work. Disease is assumed to occur at the time of 20% initial beta cell mass, which corresponds to the onset of stage 3 of the disease as discussed in Section 1. That is, we define clinical disease onset as the loss of 80% of initial beta cell mass [40], i.e. a critical time *t*_*c*_ such that

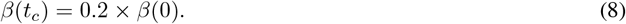

Note that if *t*_*c*_ exists (see Section 2.1), it is unique, since *β* is decreasing. We assume a non-diseased state if the critical mass of 20% of the initial beta cell mass is not reached within 70 years (i.e. we do not run our simulations longer than 70 years, as we are assuming that this approximately captures the average human lifespan [41]; mathematically this corresponds to bounding *t*_*c*_ above by 70 years, even if it is longer or does not exist). We use the T1DI model to better understand immune dynamics during the progression of type 1 diabetes, as well the effect of Treg therapy in different (patient-specific) parameter regimes.

### 2.1 Dynamics of the T1DI model

We are interested in understanding the impact of initial conditions and parameters on the time *t*_*c*_ to stage 3 disease (clinical onset defined by equation (8)). We begin by exploring the possible dynamics of the T1DI model with no external Treg therapy (*U* (*t*) ≡ 0). Define

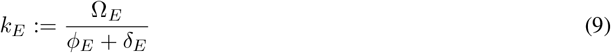

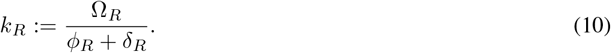

Biologically, *k*_*E*_ and *k*_*R*_ represent the *relative* growth rates of the effector and regulatory *T* cells in the lymph nodes, respectively. Here relative means with respect to removal (death plus transition to the pancreas), as (for example) *ϕ*_*E*_ + *δ*_*E*_ is the net transition rate of the effector cells out of the lymph nodes, and Ω_*E*_ denotes the maximal effector cell growth rate.

Defining *x*(*t*) := (*A*_*L*_(*t*), *E*_*L*_(*t*), *R*_*L*_(*t*), *A*_*P*_ (*t*), *E*_*P*_ (*t*), *R*_*P*_ (*t*), *β*(*t*)) ∈ ℝ^7^ as the state of the system at time *t* ≥ 0 and assuming positive parameter values and non-negative initial conditions, we can classify the asymptotic behavior of *x* into three cases. When *k*_*R*_ *> k*_*E*_,

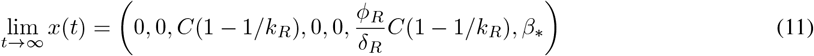

where *β*_∗_ ∈ [0, *β*(0)) is determined from the initial conditions and parameters. In this case, we say that Tregs “win” with respect to effector T cells, leading the effector T cell population in the lymph nodes and pancreas to asymptotically vanish; conversely, the Treg populations persist. Note that the restricted (*E*_*L*_, *R*_*L*_) dynamics are essentially a classical competition model, which motivates the usage of the term “win.” In this parameter regime, Tregs are able to overcome the effector T cells, thus allowing the population of beta cells (*β*) the chance to remain positive and avoid disease; the latter is not guaranteed, and again depends on specific parameter values (e.g. not just *k*_*R*_ and *k*_*E*_) and initial conditions. Having *k*_*R*_ *> k*_*E*_ can thus be interpreted biologically as having Tregs (*R*_*L*_ and *R*_*P*_) be more fit compared with the effector T cell populations (*E*_*L*_ and *E*_*P*_).

The case *k*_*R*_ *< k*_*E*_ is similar, in that

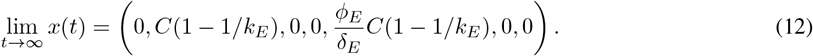

Here, we observe essentially the opposite of the previous scenario: as *t* → ∞, Tregs in the lymph node and pancreas vanish while the effector T cell populations persist. In this case, effector T cells appear to “win”, leading to a beta cell mass approaching 0 as *t* → ∞. Thus, effector T cells may be classified as more fit than Tregs.

When *k*_*R*_ = *k*_*E*_, we obtain

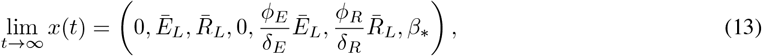

where *Ē*_*L*_ and 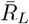 lie on the line 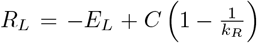, and again *β*_∗_ ∈ [0, *β*(0)). In this case we obtain an asymptotic coexistence between Treg and effector T cell populations, with both being equally “fit;” the asymptotic population sizes *Ē*_*L*_ and 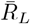 are determined by initial conditions, with *R*_*L*_(0) inversely related to *Ē*_*L*_ (see Figure 1). Intuitively, we see that when this coexistence is such that *Ē*_*L*_ is near zero on the above line, Tregs are favored; when coexistence is such that 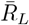 is near zero, effector T cells are favored. Thus, as the disease progresses, neither Tregs or effector T cells are able to fully replace the other, and beta cell dynamics (and hence T1D progression) are highly dependent on the degree of coexistence. This is most intuitively understood via equation (7), which for large *t* can be approximately written as

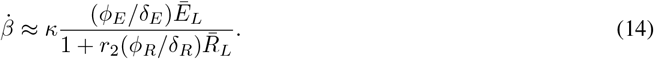

**Figure 1.**
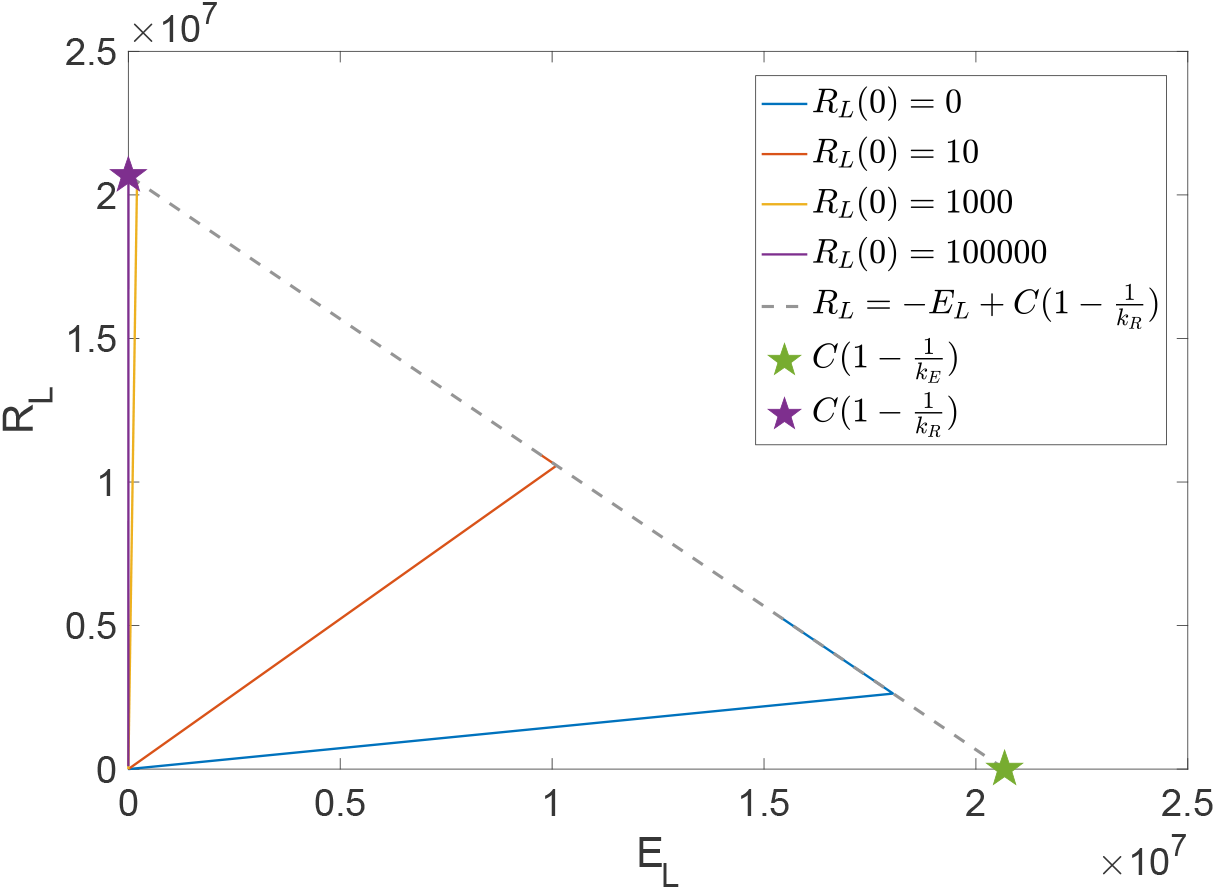
Projected dynamics of system (1) - (7) into the (*E*_*L*_, *R*_*L*_) plane for the *k*_*E*_ = *k*_*R*_ case. Here *E*_*L*_(0) = 10, *R*_*L*_(0) varies as specified in the legend, *B*(0) = 1 × 10^6^, and all other initial conditions are 0. Treg therapy is not applied, so that *U* ≡ (*t*) 0. We fixed *r*_1_ = *r*_2_ = 1 and *ν* = 1 × 10^−5^ arbitrarily, and all other parameter values are as specified in Table 2.

From the above, we directly observe the effect of *Ē*_*L*_ and 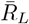 on the time *t*_*c*_ to clinical disease onset; this approximation is of course only asymptotic, as the convergence rate is also important in precisely understanding beta cell decay.

In the following sections, we attempt to understand the case when *k*_*R*_ = *k*_*E*_ in the context of quantity-versus-quality of Tregs in disease dynamics. We focus on the latter *k*_*R*_ = *k*_*E*_ case, as although there is a known imbalance in effector and regulatory T cells in individuals with type 1 diabetes, we do not expect a complete elimination of either immune component [42], which we know must occur if either *k*_*R*_ *> k*_*E*_ or *k*_*R*_ *< k*_*E*_. Furthermore, by standard continuity arguments for parameterized ODEs, our results will not change qualitatively for small parameter variations [43]. Proofs of all statements in this section are provided in Section 4.7.

### 2.2 How regulatory T cell *quality* impacts timing of disease onset

We begin by analyzing the impact of the two Treg mechanisms on disease onset by isolating each mechanism individually. The first Treg mechanism, controlled by the parameter *r*_1_, occurs in the lymph node compartment and functions by suppressing the activation of effector T cells. The second Treg mechanism, controlled by the parameter *r*_2_, directly suppresses the apoptotic capabilities of Tregs in the pancreatic compartment.

We initially fix *r*_2_ = 0 and examine the impact on the timing of disease onset *t*_*c*_ to variations in *r*_1_ and *ν*. In particular, we investigate *ν* due to the wide range of values reported in the literature (see Table 2). Furthermore, *ν* is a function of the parameter *δ*_*P*_ (decay rate of peptides in the pancreas; see equation (32) in Section 4.1.2 a model which includes peptide dynamics) which has been shown to be important in disease progression [24, 25]. In Figure 2a we vary *ν* values in ranges provided in Table 2 and vary *r*_1_ to scale *R*_*L*_; that is, we scale *r*_1_ so that *R*_*L*_ does not immediately suppress the APC activation of *E*_*L*_ in equation (26). Figure 2a provides the time to disease onset *t*_*c*_ when varying both *r*_1_ and *ν*. Interestingly, we observe two distinct parameter regimes. First, for small *ν*, increasing *r*_1_ does not protect against early disease onset (*t*_*c*_ *<* 10 years for all *r*_1_ when *ν* is approximately smaller than 10^−5^). On the other hand, for larger *ν*, increasing *r*_1_ directly results in an increase in time to disease. This suggests that improving the quality of Treg-based suppression of effector T cell activation in the lymph nodes, in the absence of any suppressive activity in the pancreas, can significantly prolong disease onset for large *ν*. It however has no effect on disease onset in the small *ν* case.

**Figure 2.**
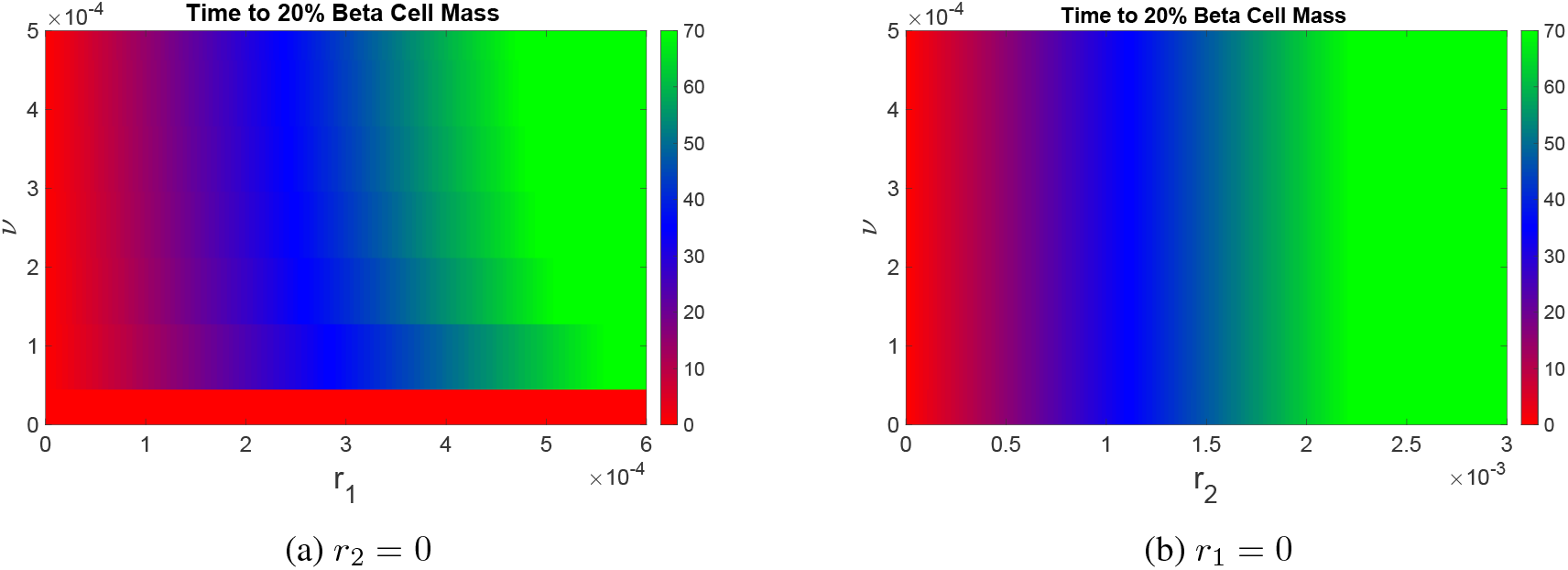
Time *t*_*c*_ to 20% beta cell mass of the T1DI model when varying (a) *ν* and *r*_1_ while setting *r*_2_ = 0 and (b) *ν* and *r*_2_ while setting *r*_1_ = 0. Initial conditions utilized are *R*_*L*_(0) = *E*_*L*_(0) = 10 and *B*(0) = 1 × 10^6^, while other initial populations are 0. All other parameters may be found in Table 2. Treg therapy is not applied, so that *U* (*t*) ≡ 0. The discretization of *r*_1_ and *ν* occurs on a mesh grid of size 300 x 300 (see Figure 9 for a higher resolution parameter discretization). Red indicates early disease onset, blue indicates late disease onset, and green indicates no disease onset.

We next perform an analogous experiment, in which we vary *ν* and *r*_2_ while fixing *r*_1_ = 0. Results are provided in Figure 2b, where we observe no distinction between a small and large *ν* regime. Instead, we note that as *r*_2_ increases, the time to disease onset increases, with an only marginal effect due to the *ν* parameter. This is intuitive, as the *r*_2_ mechanism directly impacts the killing of beta cells by effector T cells, so that increasing *r*_2_ will directly decrease the rate at which beta cell mass is lost.

Biologically, for individuals with type 1 diabetes, it is reasonable to assume that both Treg mechanisms are functioning to some (possibly impaired) degree. Hence, we analyze disease progression when both *r*_1_ and *r*_2_ are non-zero, i.e both Treg mechanisms are at least partially functional. We vary *r*_1_ and *r*_2_ as in Figures 2a and 2b, respectively, and then fix two values of *ν* corresponding to the distinct parameter regimes as observed in Figure 2a. Figure 3b shows the time to disease onset when fixing *ν* in the large regime (*ν* = 0.01) and varying *r*_1_ and *r*_2_. Here, we see again that there exist distinct regions of early (red), late (blue), and no disease (green) onset. On the other hand, Figure 3a shows the time to disease onset when when *ν* is in the small regime (*ν* = 1 × 10^−5^). Here, we again see the three distinct disease onset times. However, we note that, when compared with Figure 3b, higher values for both *r*_1_ and *r*_2_ are necessary to achieve a no disease state. Hence, in the case of small *ν*, we conclude that greater Treg function is necessary to prevent disease.

**Figure 3.**
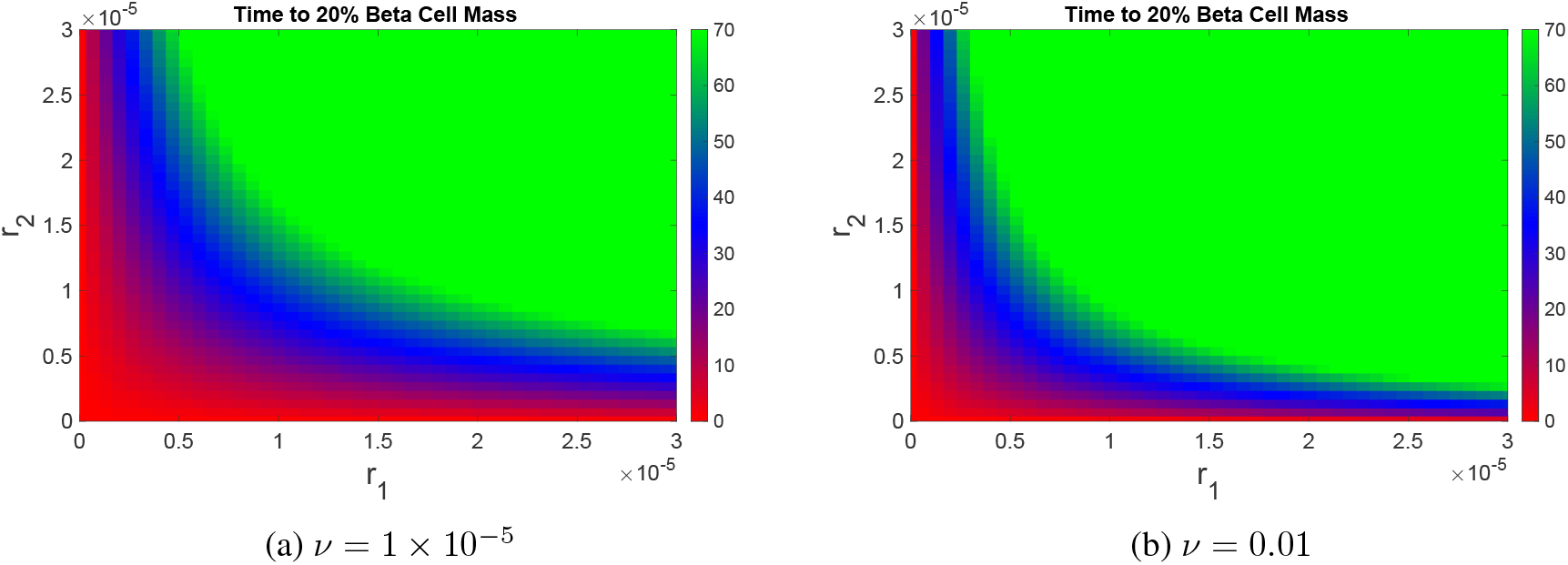
Time *t*_*c*_ to 20% beta cell mass of the T1DI model when varying *r*_1_ and *r*_2_ for (a) *ν* = 1 × 10^−5^ and (b) *ν* = 0.01. Initial conditions utilized are *R*_*L*_(0) = *E*_*L*_(0) = 10 and *B*(0) = 1 × 10^6^, while other initial populations are 0. All other parameters may be found in Table 2. Treg therapy is not applied, so that *U* (*t*) ≡ 0. The discretization of *r*_1_ and *r*_2_ occurs on a mesh grid of size 300 x 300. Red indicates early disease onset, blue indicates late disease onset, green indicates no disease onset. For any fixed pair of *r*_1_ and *r*_2_, the time to disease onset is greater for the larger *ν* parameter value (*t*_*c*_ in (b) is larger than the corresponding *t*_*c*_ in (a)).

### 2.3 How regulatory T cell *quantity* impacts timing of disease onset

In the previous section we investigated the impact of *quality* of Tregs on time *t*_*c*_ to stage 3 disease; recall that quality was quantified via parameters *r*_1_ and *r*_2_ in our T1DI model. In this section, we investigate the role of the *quantity* of regulatory T cells on *t*_*c*_. Previously, we observed that different *ν* values lead to different disease progression dynamics. Thus, in this section we also investigate the effect of *ν* in response to Treg therapy, which is modeled as an external input *U* (*t*) in equation (3). We consider a simplified model Treg therapy, where we introduce Tregs at different doses *D* and time points *t*_*D*_ in the time course of disease; we subsequently observe the resulting time to disease onset *t*_*c*_ when compared with no Treg therapy. Note that the dose time points *t*_*D*_ discussed in this section can also be translated to remaining beta cell mass via C-peptide levels, which may be easier to estimate than the time since onset of stage 1; this translation is useful since the time of stage 1 disease onset is often unknown. That is, we can also use a patient-calibrated model to determine dosing time *t*_*D*_ as a function of clinically measured beta cell mass. For example, see Table 2 in [44] for a relationship between diabetes staging and C-peptide levels. For details on the implementation of the Treg therapy dosing function *U* (*t*), see Section 4.1.3 and specifically equation (34).

As in Figure 3, we fix *ν* = 1 × 10^−5^ (small) and *ν* = 0.01 (large). We choose *r*_1_ and *r*_2_ values corresponding to both early and late disease onset found in Figures 3a and 3b. In fact, when making direct comparisons, we choose the same *r*_1_ and *r*_2_ values for both large and small *ν*, so that differences in Treg function (quality) can be excluded as a possible factor contributing to differences in response between the two *ν* values. Moreover, we quantify the change in response to Treg therapy as the difference in time to 20% beta cell mass between zero (*D* = 0) and positive (*D >* 0) dose *D*:

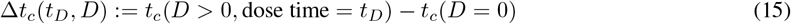

Note that additional regulatory T cells can only improve response (i.e. increase *t*_*c*_), so that Δ*t*_*c*_ is always non-negative. Larger values of Δ*t*_*c*_ thus correspond to an improved response.

Figure 4 shows the difference in time to 20% beta cell mass (Δ*t*_*c*_) when varying both the dose amount *D* and dose time *t*_*D*_ for early (*r*_1_ = 0.2 × 10^−5^ and *r*_2_ = 0.2 × 10^−5^; Figure 4a) and late (*r*_1_ = 0.8 × 10^−5^ and *r*_2_ = 1 × 10^−5^; Figure 4b) disease parameters when *ν* is small (compare to Figure 3a). In both cases, we observe that dosing Tregs is effective when dosed high enough and at the proper time; we note that we tested lower doses and found they have little to no impact on disease timing (not shown). In particular, we observe an optimal dosing window between 1 and 2 years in the early disease onset parameter case (Figure 4a) and that for high Treg doses (*>* 10^8^ cells) administered during this time, the time to 20% beta cell mass is extended by 2-4 years (Figure 4a). In the late disease parameter case (Figure 4b), Treg doses greater than 10^9^ cells result in delays of disease onset by more than 15 years at any dosing at any time. The optimal results are obtained when cells are administered within 15 - 30 years, where time to 20% beta cell mass is extended by as much as 40 years, which yields complete disease prevention. Hence, we find for both cases that there is an optimal *intermediate* dosing window. That is, for the largest increase in time to disease onset, Tregs must not be applied too early nor too late. However, for late disease onset parameters, even dosing outside of the optimal window results in a large delay of disease, i.e. the response is much more robust with respect to proper timing. We note that Teplizumab (brand name Tzield®), an FDA approved drug to delay the onset of T1D, is dosed during stage 2 of disease progression (i.e. during an intermediate stage of disease progression) [45].

**Figure 4.**
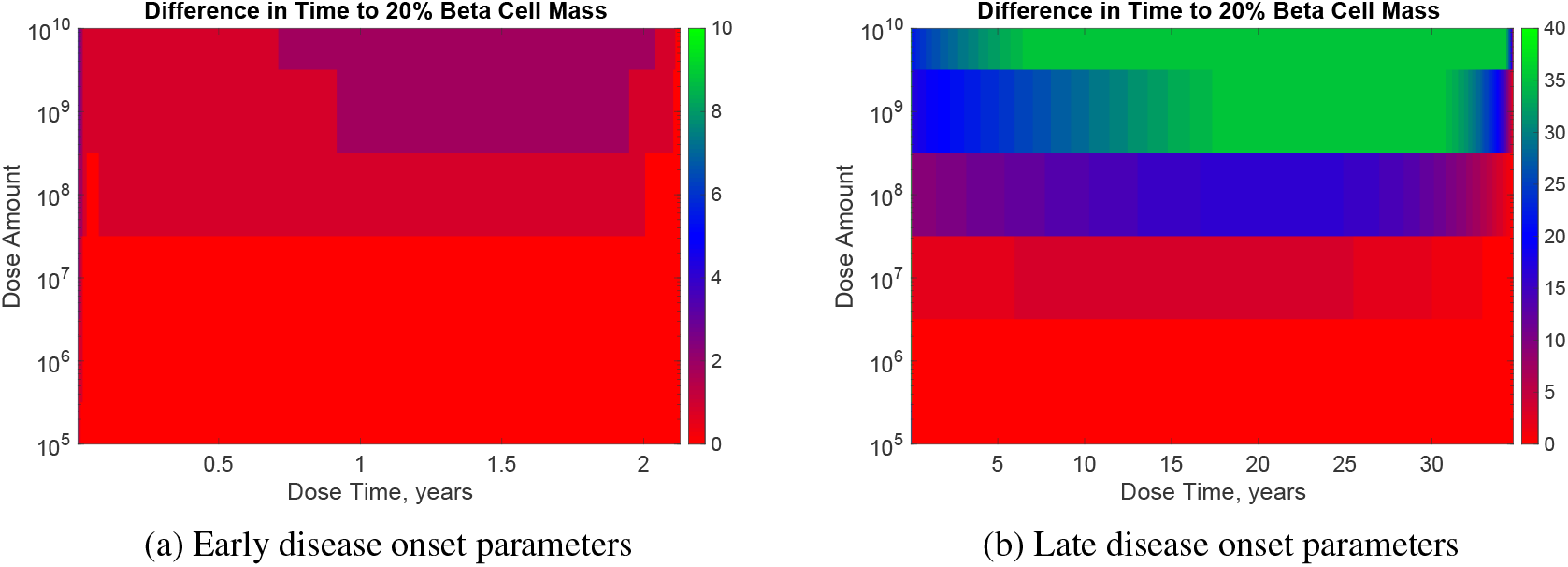
Response to Treg therapy in small *ν* parameter regime. Increases in time to 20% beta cell mass (Δ*t*_*c*_ in (15)) when varying *t*_*D*_ and *D* in regulatory T cell therapy for *ν* = 1 × 10^−5^ for (a) early disease onset (*r*_1_ = 0.2 × 10^−5^ and *r*_2_ = 0.2 × 10^−5^) and (b) late disease onset (*r*_1_ = 0.8 × 10^−5^ and *r*_2_ = 1× 10^−5^) parameters. Initial conditions utilized are *R*_*L*_(0) = *E*_*L*_(0) = 10 and *B*(0) = 1 × 10^6^, while other initial populations are 0. All other parameters may be found in Table 2. Relative disease onset time Δ*t*_*c*_ is plotted as a function of both *t*_*D*_ (*x*-axis) and *D* (*y*-axis). The discretization of *t*_*D*_ and *D* occurs on a mesh grid of size 300 x 300. Red indicates no change in time to 20% beta cell mass when compared with not dosing (i.e. no response), blue indicates a slight increase in time to 20% beta cell mass, and green indicates the largest increase in time to 20% beta cell mass (i.e. maximal response).

To understand the intermediate optimal dosing window observed in Figure 4, we investigate the relationship between *E*_*P*_ × *β* at the dose time *t*_*D*_ and the time to 20% beta cell mass *t*_*c*_; Figure 5 plots this relationship for the late disease onset parameters (Figure 4b). Here, we observe a clear pattern: small *E*_*P*_ × *β* at the time of dose produces a large time to 20% beta cell mass. Thus dosing at smaller values of *E*_*P*_ × *β* yields an improved disease outcome. We note that this phenomenon does not appear to be true when the dose time is very close to disease onset, in which case Treg dosing has little effect (see Figure 11). This latter lack of increased *t*_*c*_ is not surprising, as the inherent delays in the T1DI model do not allow for instantaneous responses.

**Figure 5.**
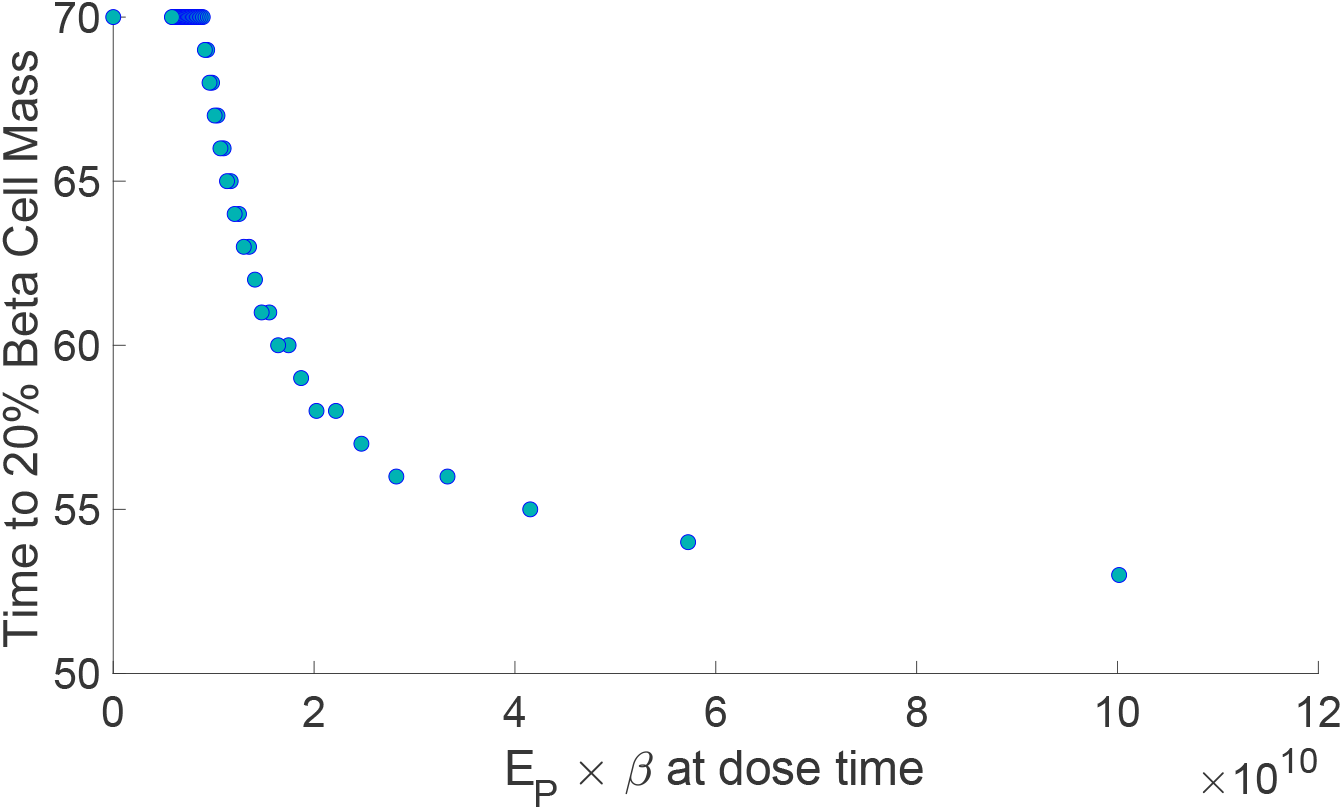
Time to 20% beta cell mass *t*_*c*_ versus *E*_*P*_ × *β* at dose time *t*_*D*_. Here, we fixed parameters as in Figure 4b, i.e. we consider late disease onset parameters (*r*_1_ = 0.8 × 10^−5^ and *r*_2_ = 1 × 10^−5^) for *ν* = 1 × 10^−5^. We fix the dose as *D* = 1 × 10^9^ cells. Initial conditions utilized are *R*_*L*_(0) = *E*_*L*_(0) = 10 and *B*(0) = 1 × 10^6^, while other initial populations are 0. All other parameters may be found in Table 2.

Mechanistically, we hypothesize that *t*_*c*_ will decrease as *E*_*P*_ × *β* increases, since *E*_*P*_ × *β* is proportional to the rate of APC activation (see equation (4); APCs are produced when effector T cells and beta cells interact in the pancreas at a rate *νE*_*p*_*β/*(1 + *r*_2_*R*_*p*_)). When APC activation is small, the dynamics of effector and regulatory T cells in the lymph nodes are driven by competition between the two populations (carrying capacity *C* in equations (2) and (3)), and an external dose of Tregs *U* will “tip the balance” in the favor of regulatory T cells, thus suppressing the autoimmune response and increasing *t*_*c*_. Analogously, when APC activation is large (large *E*_*P*_ × *β*), the dynamics of the immune system in the lymph nodes are driven by APC (*A*_*L*_), with little additional effect due to *U*; thus, we expect to observe a small change in *t*_*c*_ (Δ*t*_*c*_ ≈ 0). See Section 4.6 for further analysis and discussion of the effect of APC on dynamic response in a simplified model of immune competition.

We also tested the difference in time to 20% beta cell mass when varying both the dose amount and dose time for early and late disease parameters for large *ν* (compare to Figure 3b). However, contrary to the small *ν* case (Figure 4), here we find that for both early and late disease onset parameters there is negligible improvement in response to Treg dosing regardless of dose time or amount (the maximum Δ*t*_*c*_ for both early and late disease is on the order of 10^−5^ years), and hence a figure analogous to Figure 4 is not included. This is biologically important as it suggests that a difference in response to Treg therapy depends on a patient’s *ν* value: individuals with small *ν* will exhibit a significant delay in disease onset, while those with a larger *ν* will not.

As mentioned previously, we hypothesize that the difference in response to Treg therapy between small and large *ν* may be due to APC dynamics, which we have investigated in Section 4.6 with two reduced models. Specifically, we analyze a model (equations (44) - (45)) in which the number of APC cells is a fixed parameter *a* ≥ 0. Simulating, we find that varying the constant amount of APC yielded similar dynamics as observed when varying *ν* (Figures 12a and 12b, respectively). That is, we see that smaller *a* corresponds to a positive response to Treg therapy (increased *t*_*c*_), which is not observed for larger *a*. These results can be understood analytically via perturbation theory/phase plane analysis of a further reduced model (46) - (48), which is also included in Section 4.6. Hence, we conclude that APC dynamics may be driving the response to Treg therapy. This difference in “disease driver” leads us to investigate the dynamics of the T1DI model when we deplete APCs at the time of Treg dosing.

We note that we have also performed a sensitivity analysis on all model parameters except *r*_1_, *r*_2_, and *ν*, as these latter parameters are systematically investigated in this work. See Section 4.4, as well as Figure 10, for details, which are briefly summarized here. Sensitivity is computed with respect to time *t*_*c*_ to 20% initial beta cell mass, and is measured as the slope of *t*_*c*_ with respect to fold changes in the corresponding parameter. We determined that *δ*_*E*_ and *δ*_*R*_ were among the most sensitive parameters, with *t*_*c*_ increasing rapidly with respect to *δ*_*E*_, while time to disease decreases with respect to *δ*_*R*_. Furthermore, we also observe that the system is sensitive to *λ*_*E*_ and *λ*_*R*_: *t*_*c*_ decreases with respect to *λ*_*E*_, and increases with respect to *λ*_*R*_. These parameters further suggest that the relative fitness of regulatory versus effector T cells is important to the dynamics of the T1DI model, as the sensitive parameters are all related to (*E*_*L*_, *R*_*L*_) interactions.

### 2.4 Depleting APC at time of regulatory T cell dose improves response

The depletion of APCs (primarily B cells) has been studied in autoimmune diseases such as lupus and multiple sclerosis [46, 47]; it has also been studied in T1D [48, 49]. A phase I/II trial in patients with new-onset T1D found that patients who received a combination of rituximab, which binds to CD20 on B cells and can elicit apoptosis, together with Treg therapy exhibited an improved response compared to those who received rituximab or Treg therapy alone [48]. In this section, we explore an idealized scenario of combination therapy in which we dose Tregs and deplete APCs simultaneously. To model the depletion of APCs, we fully suppress their function, i.e. we remove their ability to activate effector T cells or Tregs in the lymph nodes. See Section 4.1.4 (specifically equations (42) and (43)) for precise details of the implementation of APC deletion in the T1DI model (1) - (7).

Figure 6 shows the difference in time to 20% beta cell mass Δ*t*_*c*_ when varying the dose amount *D* and dose time *t*_*D*_ for Tregs while simultaneously depleting APCs for small *ν* (*ν* = 1 × 10^−5^). Here, we observe that for both early and late disease parameters, the response improves when compared to only dosing Tregs without depleting APCs (Figure 4). In fact, for larger doses *D*, total remission (i.e. stage 3 of the disease is not obtained within 70 years) occurs. Moreover, compared with monotherapy, the optimal dosing time *t*_*D*_ occurs earlier. Similar results hold for late disease parameters, which are provided in Figure 6b. Results for an analogous simulation for the high *ν* case are provided in Figure 7. When compared with the Treg monotheray, which exhibited essentially no improvement in time to disease, we observe that the large *ν* regime responds significantly to the combination therapy, with complete prevention possible for sufficiently high doses. Furthermore, dosing earlier appears to be optimal, and that for late disease onset parameters (Figure 7b), depleting APC early (but not too early; approximately 10 years) is sufficient to delay disease onset, even at low doses of Tregs.

**Figure 6.**
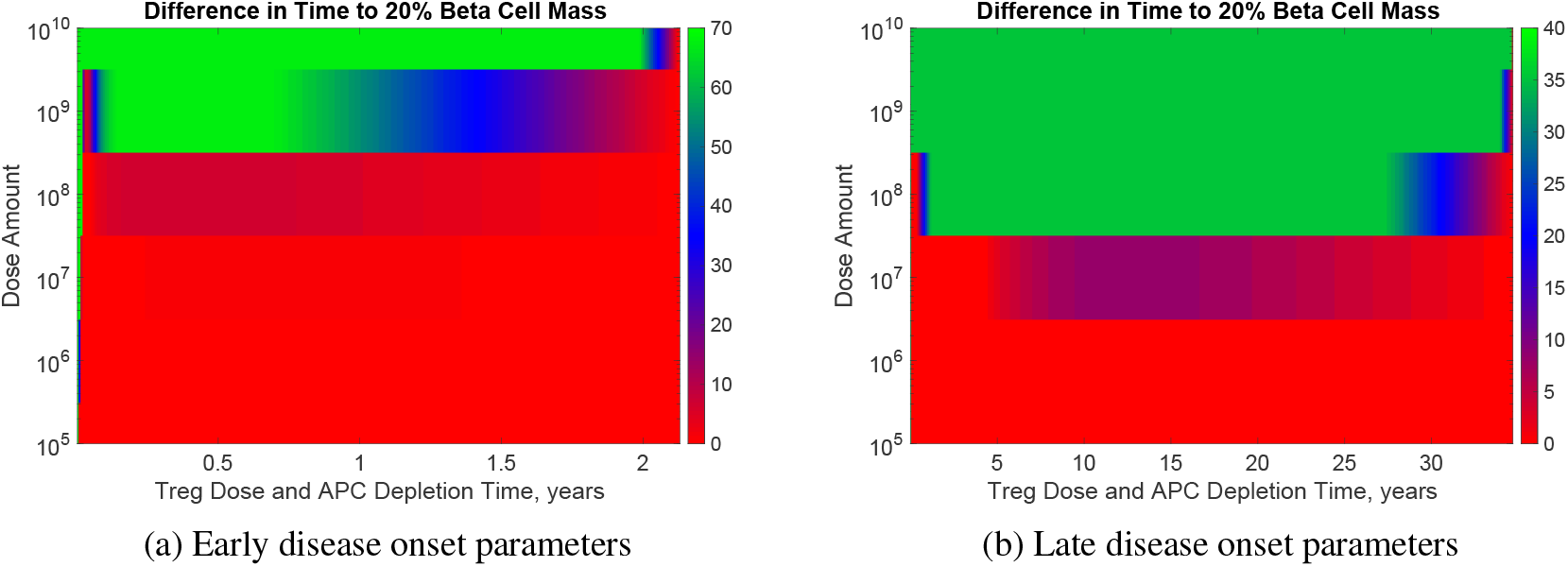
Time to 20% beta cell mass when varying Treg dose *D* and dose time *t*_*D*_ in combination with APC depletion for small *ν*. Increases in time to 20% beta cell mass (Δ*t*_*c*_ in (15)) when varying *t*_*D*_ and *D* in regulatory T cell therapy for *ν* = 1 × 10^−5^ for (a) early disease onset (*r*_1_ = 0.2 × 10^−5^ and *r*_2_ = 0.2 × 10^−5^) and (b) late disease onset (*r*_1_ = 0.8 × 10^−5^ and *r*_2_ = 1 × 10^−5^) parameters. Initial conditions utilized are *R*_*L*_(0) = *E*_*L*_(0) = 10 and *B*(0) = 1 × 10^6^, while other initial populations are 0. All other parameters may be found in Table 2. Relative disease onset time Δ*t*_*c*_ is plotted as a function of both *t*_*D*_ (*x*-axis) and *D* (*y*-axis). The discretization of *t*_*D*_ and *D* occurs on a mesh grid of size 300 x 300. Red indicates no change in time to 20% beta cell mass when compared with not dosing (i.e. no response), blue indicates a slight increase in time to 20% beta cell mass, and green indicates the largest increase in time to 20% beta cell mass (i.e. maximal response). Recall that APCs are depleted at dose time *t*_*D*_.

**Figure 7.**
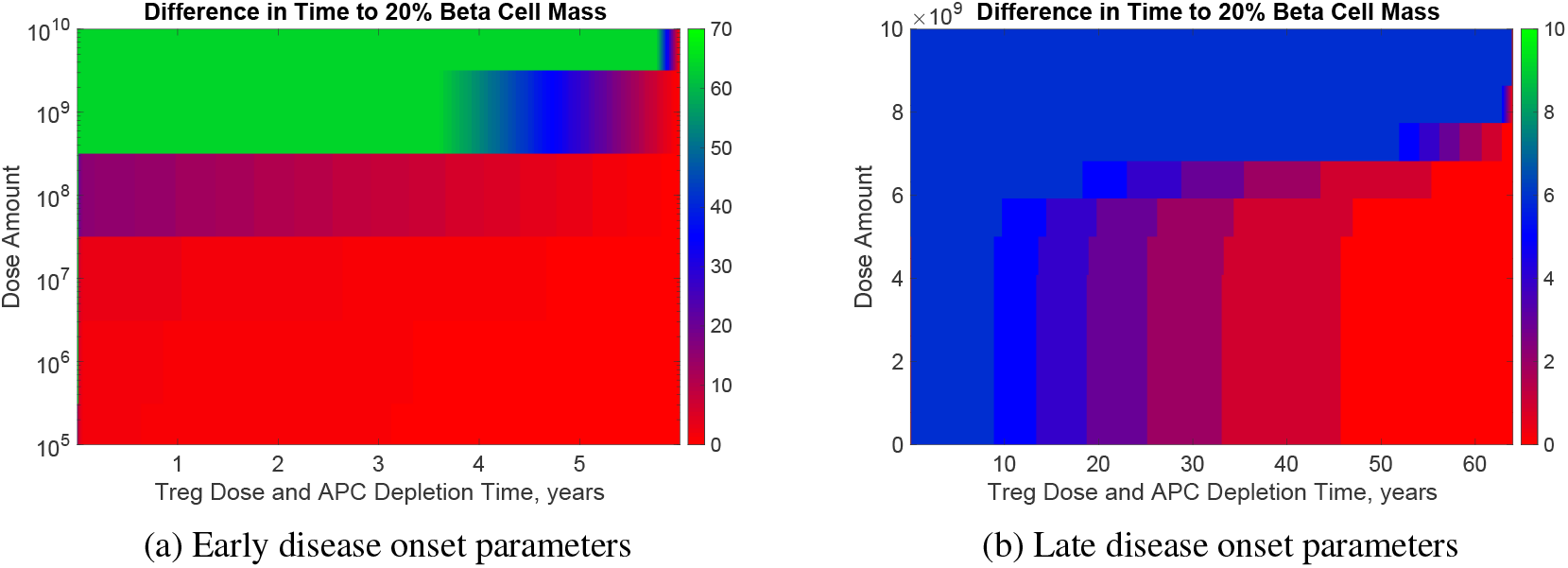
Time to 20% beta cell mass when varying Treg dose *D* and dose time *t*_*D*_ in combination with APC depletion for high *ν*. Increases in time to 20% beta cell mass (Δ*t*_*c*_ in (15)) when varying *t*_*D*_ and *D* in regulatory T cell therapy for *ν* = 0.01 for (a) early disease onset (*r*_1_ = 0.2 × 10^−5^ and *r*_2_ = 0.2 × 10^−5^) and (b) late disease onset (*r*_1_ = 0.8 × 10^−5^ and *r*_2_ = 1 × 10^−5^) parameters. Initial conditions utilized are *R*_*L*_(0) = *E*_*L*_(0) = 10 and *B*(0) = 1 × 10^6^, while other initial populations are 0. All other parameters may be found in Table 2. Relative disease onset time Δ*t*_*c*_ is plotted as a function of both *t*_*D*_ (*x*-axis) and *D* (*y*-axis). The discretization of *t*_*D*_ and *D* occurs on a mesh grid of size 300 x 300. Red indicates no change in time to 20% beta cell mass when compared with not dosing (i.e. no response), blue indicates a slight increase in time to 20% beta cell mass, and green indicates the largest increase in time to 20% beta cell mass (i.e. maximal response). Recall that APCs are depleted at dose time *t*_*D*_.

## 3 Discussion and conclusions

In this work, we have developed a two compartment mathematical model to understand how the *quality* and *quantity* of regulatory T cells impact timing to clinical disease onset in type 1 diabetes. We began by analyzing the impact of regulatory T cell *quality* on time to 20% beta cell mass. By testing two regulatory mechanisms (modeled via *r*_1_ and *r*_2_ in system (1) - (7)) separately and varying *ν* in physiological ranges reported in literature, we discovered two distinct parameter regions for *ν* that resulted in qualitatively different disease dynamics. Specifically, for sufficiently small *ν*, the *r*_1_ mechanism (which models the suppression of effector T cell activation) was not protective against disease. Biologically this can be explained as follows: small *ν* implies small early APC populations, so that there is minimal effector T cell activation via APCs in the lymph nodes; recall that APC activation is inhibited by *r*_1_. In this case, the increase of effector T cells occurs mainly through proliferation. When studying the *r*_2_ mechanism (which models the suppression of apoptotic ability of effector T cells), distinct parameter regimes for *ν* were not observed.

A major finding of this work is that the parameter *ν* is important in disease progression, both in the absence and presence of Treg therapy. Recall from equation (32) that *ν* is defined via three rate parameters: the activation rate of APC by peptide (*σ*), the accumulation rate of peptide (*α*), and the clearance rate of peptide (*δ*_*P*_). A large *ν* value may thus arise via one or more of the following physiological scenarios: a high rate of activation of APC by peptide, a large accumulation rate of peptide, and/or a slow clearance of peptide. We note that it has been observed that small increases in peptide clearance rates (*δ*_*P*_ in our model) result in a healthy state [25]. Similarly, it has also been observed that changes in macrophage clearance rates (which is analogous to *δ*_*P*_ in our model) altered the trajectory of disease [24]. Specifically, it was found that high peptide clearance rates (low *ν*) yielded a non-diabetic mouse, whereas low clearance levels (high *ν*) produced high glucose levels, i.e. a diabetic mouse. Hence, we speculate that the parameter *δ*_*P*_ may lead to the differences in disease progression that we have demonstrated in our analysis of parameter *ν*.

Once we discovered the two distinct parameter regimes for *ν*, we investigated variations in both regulatory mechanisms *r*_1_ and *r*_2_ simultaneously, as it is biological reasonable to assume both mechanisms are at least partially functional. We found that time to disease onset was more sensitive to changes in *r*_1_ and *r*_2_ in the case of case of small *ν* (*ν* = 1 × 10^−5^), as larger values of *r*_1_ and *r*_2_ were needed to remain disease free in this case. Hence, for individuals with low peptide-induced APC activation rate, regulatory T cells must generally have increased efficacy to prevent the onset of type 1 diabetes.

After studying the quality of regulatory T cells, we analyzed regulatory T cell *quantity* and its impact on time to 20% beta cell mass. We again distinguished between small and large *ν* parameters values, and fixed quality parameters *r*_1_ and *r*_2_ that corresponded with early and late disease onset. Quantity of Tregs is directly related to the dosing regulatory T cells, so we were interested in the role of parameter *ν* and response to treatment. Through simulations and mathematical analysis, we discovered that for small *ν* values, a favorable response to regulatory T cell therapy was observed, and furthermore, that there exists an optimal dosing window. On the other hand, we found that for larger *ν*, there is no response to regulatory T cell therapy. We hypothesized that the differences in response to regulatory T cell therapy between small and large *ν* was a consequence of APC dynamics, which again was theoretically justified via mathematical models. Hence, we modeled the depletion of APCs at the time of regulatory T cell dosing, and studied the impact of this combination therapy on time to 20% beta cell mass. Through simulations, a significant response to therapy in both the small and large *ν* parameter regimes was observed, and that the response to dosing is more robust when compared to the case of Treg monotherapy. Indeed, in some cases, remaining disease free for more than 70 years was possible when applying combination therapy.

Using mechanistic modeling, we have investigated the role of regulatory T cell quality and quantity, and how they can be related to response to Treg therapy, a current avenue of research in the treatment of type 1 diabetes. We have shown that certain physiological parameters play a key role in treatment response, and have shown that combining Treg therapy with APC depletion therapy may be a possible avenue to improve efficacy. Beyond applications to type 1 diabetes, our modeling framework may have applications in autoimmune diseases more generally, and future work is needed to both calibrate models to in vitro and in vivo data, as well as to incorporate both pharmacokinetics and pharmacodynamics into our type 1 diabetes immune model, especially in regards to APC depletion therapy. Nevertheless, the model presented provides valuable initial insights into immune dynamics leading to disease.

### Limitations of the study

As in any modeling study, a fundamental limitation is the biological simplifications necessary to construct mathematical descriptions that remain analytically tractable. We have indeed substantially simplified the role of the immune system in type 1 diabetes, but nevertheless believe the model presented includes enough detail to address the questions of interest related to regulatory T cell function.

Another limitation is related to parameter estimates utilized in simulations. APC parameter estimates relied mainly on parameter values from [19] where parameter values from professional APCs (pAPC) were used, which include dendritic cells, macrophages, and B cells. The exception to this is the trafficking rate of APC from tissue to lymph nodes (*ϕ*_*A*_), where we considered values from dendritic cells found in [50]. However, we note that we do not expect this parameter to have significant impact on our model behavior (see the sensitivity analysis for *ϕ*_*A*_ provided in Figure 10).

Lastly, we idealized APC depletion in Section 2.4 when considering combination therapy. Specifically, we assumed that APC function was completely suppressed at the time *t*_*D*_ of therapy, which is of course unrealistic (see equations (42) - (43)). We believe this idealization is justified, as our goal was not to mechanistically model combination therapy, but instead provide evidence for the role of APCs in Treg therapy efficacy. In future work, we will investigate more realistic models, including analyzing the response to the timing of APC depletion relative to when Tregs are dosed, as well as what percent of APCs need to be depleted to completely avoid disease. We believe the simplification taken in this work is sufficient as an initial step towards simply elucidating the role of APC dynamics in Treg therapy.

## Data and code availability

Code for all figures and simulations may be found at the following GitHub address: https://github.com/kenziealyse/ImmunePaper

## Author contributions

M.D., E.A.A., and J.M.G. conceptualized and supervised the study, designed the model, and developed the computational framework. M.D. performed simulations, parameter estimation from literature, sensitivity analyses, and most other numerical computations. All authors wrote and approved the manuscript.

## 4 Methods

### 4.1 Mathematical models

We construct a number of mathematical models to study the role of the immune system in type 1 diabetes (T1D). The biological assumptions underlying each model are discussed at length in the subsequent subsections.

#### 4.1.1 Initial type 1 diabetes immune model

We begin by formulating a two compartment model of the immune system interacting with the pancreatic beta cells that includes antigen-presenting cells (APCs), regulatory T cells (Tregs), effector T cells, peptide, and beta cell dynamics. A schematic of the primary interactions considered is provided in Figure 8. The model consists of both lymph node and pancreatic compartments. We consider multiple compartments to capture two distinct Treg suppression mechanisms, as discussed below.

**Figure 8.**
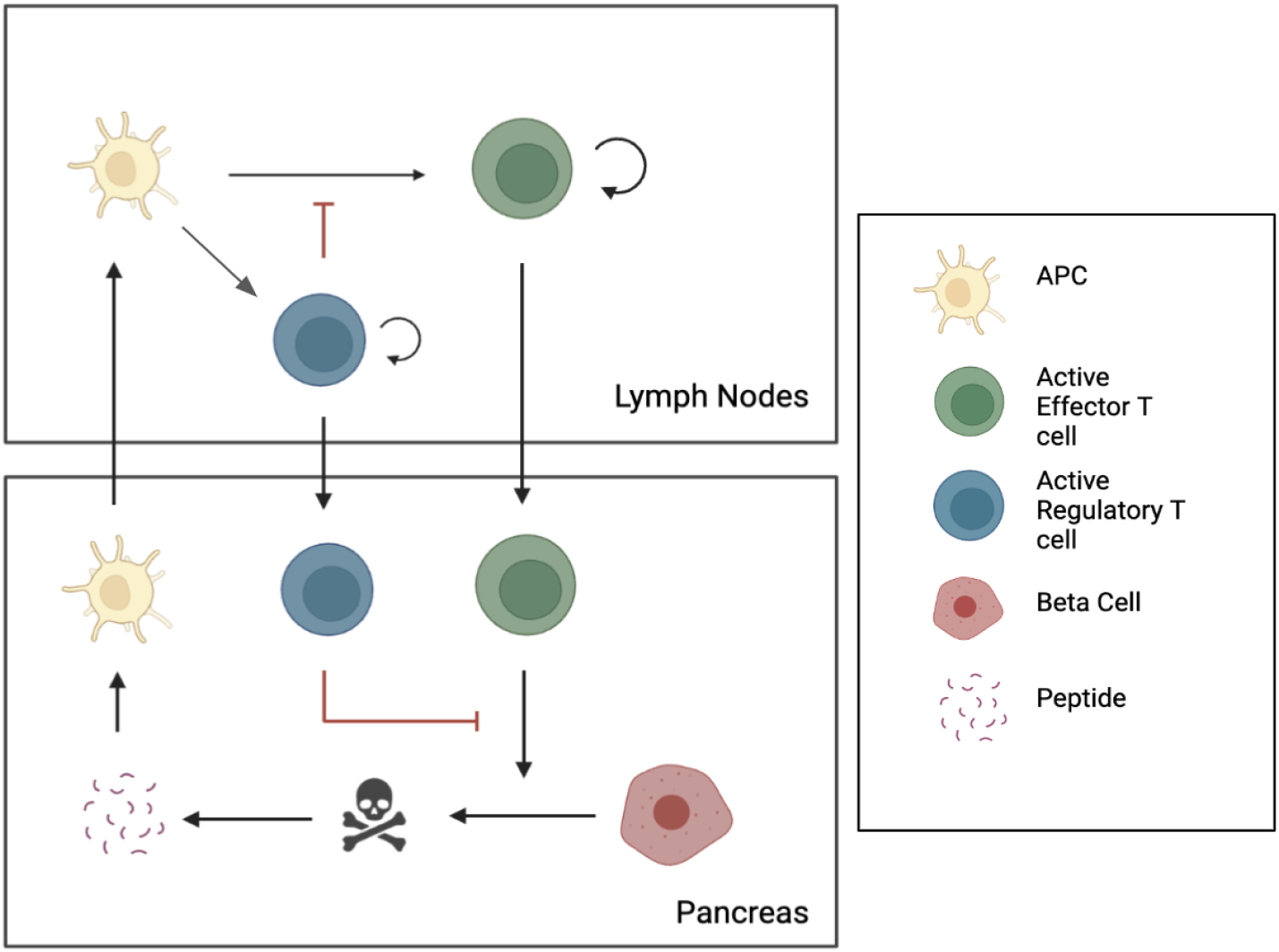
Model schematic for system (16) - (23), which describes immune and beta cell interactions in the lymph nodes and pancreas.

In the lymph nodes, APCs (*A*_*L*_), Tregs (*R*_*L*_), and effector T cells (*E*_*L*_) interact. APCs activate both regulatory and effector T cells. Tregs are able to suppress the APC activation of effector T cells, and activated Tregs and effector T cells are able to proliferate, where we assume they compete for resources. Both regulatory and effector T cells may travel to the pancreas, where APCs (*A*_*P*_), peptide, (*P*), Tregs (*R*_*P*_), effector T cells (*E*_*P*_), and beta cells (*β*) reside. In the pancreas, effector T cells kill beta cells and Tregs suppress the killing of beta cells by effector T cells. The dead beta cells also produce peptide which in turn activates APCs, which may travel to the lymph nodes [50].

We model cell populations in the lymph node compartment with three ordinary differential equations (ODEs) as follows:

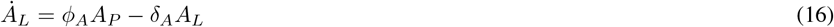

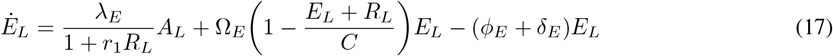

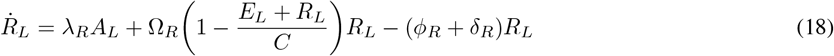

Similarly, the cells in the pancreas compartment can be described with the following five ODEs:

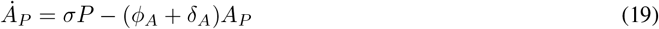

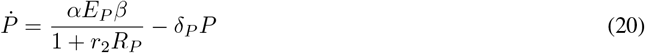

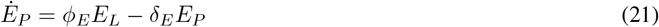

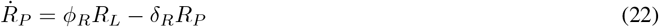

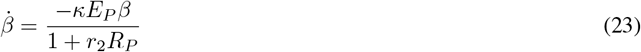

Thus, the dynamical system (16) - (23) consists of eight nonlinear ODEs. Mass action kinetics are assumed for most rate terms for simplicity. The major biological features of the model are described as follows:

1. APCs (*A*_*P*_) are activated in the pancreas through contact with self peptides (*P*) from beta cells [19]. We assume that there exists a large pool of resting APCs, and hence model the activation of APCs via *σP* (equation (19)), where *σ >* 0 is a constant.
2. In the lymph nodes, effector T cells (*E*_*L*_) are activated by APCs. As in [19], we assume a constant pool of resting effector T cells from which interaction with APCs results in an active effector T cell (*λ*_*E*_*A*_*L*_). We also model the suppression of APC activation by regulatory T cells phenomenologically as 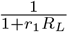 (equation (17)), so that *r*_1_ ≥ 0 controls the degree to which regulatory T cells suppressor effector cell activation by APCs in the lymph nodes.
3. In the lymph nodes, Tregs are activated by APCs. Similarly to [19], we assume that there is a constant pool of Tregs from which interaction with APCs results in active regulatory T cells (*λ*_*R*_*A*_*L*_) [51] (equation (18)).
4. In the lymph nodes, active effector T cells (*E*_*L*_) and active Tregs (*R*_*L*_) proliferate at maximal rates Ω_*E*_ and Ω_*R*_, respectively. Note that we assume no proliferation occurs in the pancreas [52]. Proliferation in the lymph nodes is inhibited by competition for resources, which is modeled via a joint carrying capacity *C*.
5. Beta cells are killed by effector T cells in the pancreas at a maximal rate of *κ* [25]. The killing of beta cells is suppressed by Tregs [53] (equation (23)), and the efficacy of this regulation is controlled by parameter *r*_2_ ≥ 0.
6. Self peptide (*P*) is produced at a rate proportional (via constant *α >* 0) to beta cell decay [25]. The production of self peptide is suppressed by Tregs due to Tregs inhibiting the killing of beta cells by effector T cells [53] (equation (20)).

Table 1 summarizes the main components of the immune-beta cell interaction model.

**Table 1:** Cell populations modeled in the immune-beta cell system (16) - (23).

| Species | Description | Initial population (cells) | Reference |
| --- | --- | --- | --- |
| $A_L$ | APCs in the lymph nodes | 0 | |
| $E_L$ | Active effector T cells in the lymph nodes | 10 | |
| $R_L$ | Active Tregs in the lymph nodes | 10 | |
| $A_P$ | APC in the pancreas | 0 | |
| $P$ | Self peptide from beta cells in the pancreas | 0 | |
| $E_P$ | Active effector T cells in the pancreas | 0 | |
| $R_P$ | Active Tregs in the pancreas | 0 | |
| $\beta$ | Beta cells | $1 \times 10^6$ | [54] |

**Table 2:**
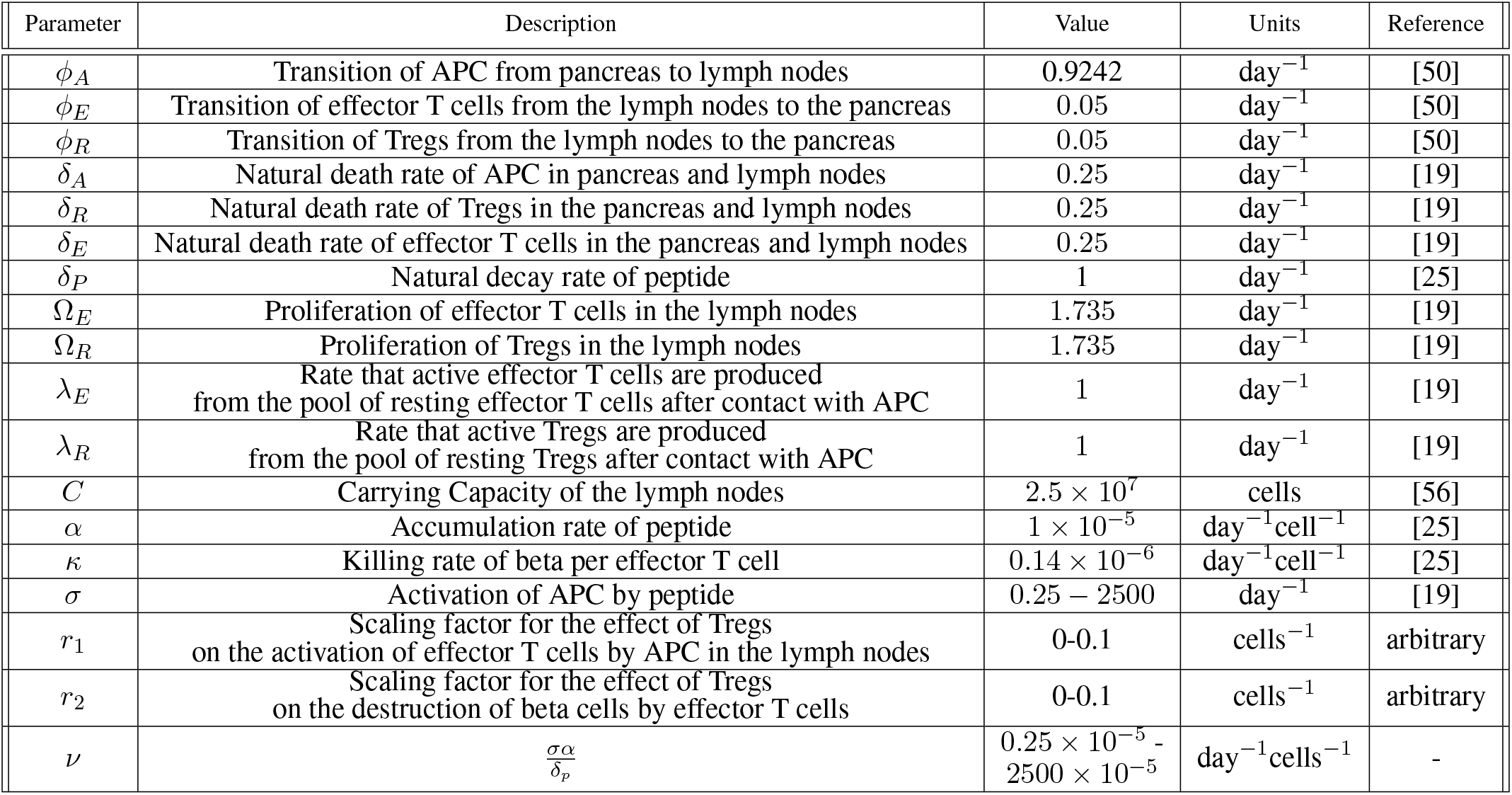
Summary of parameters utilized in simulations of the T1DI model (35) - (41).

#### 4.1.2 Quasi-steady-state approximation (T1DI model)

Motivated by natural time scale separations, we reduce the dimensionality of the immune-beta cell model (16) - (23) via a standard quasi-steady state (QSS) approximation [25]. Since the characteristic timescale of peptide dynamics is shorter than that of the other populations (hours versus days), we make a QSS approximation with respect to *P*. That is, we assume that the peptide population approaches equilibrium more rapidly than the other dynamics in equations (16) - (23). Thus we set 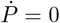 and obtain

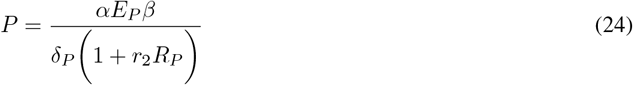

Substituting the above into equation (19), the QSS approximation results in the following reduced ODE system:

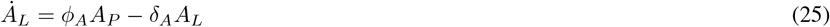

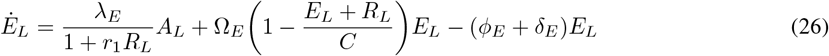

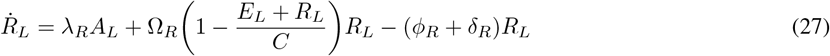

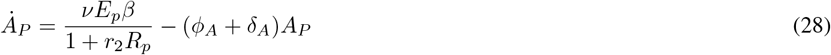

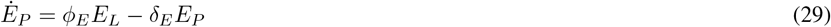

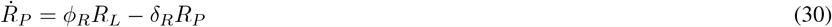

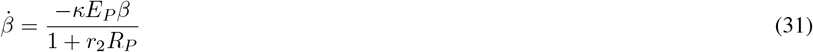

where we define the parameter *ν* as

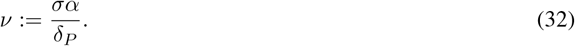

We refer to *ν* as the *peptide-induced APC activation rate*. As in the main text of the manuscript, we refer to the above model as the type 1 diabetes immune (T1DI) model; this is the primary model analyzed/simulated in this work.

#### 4.1.3 Type 1 diabetes immune model with regulatory T cell dosing

Tregs offer a potential therapeutic intervention for T1D. However, clinical trials of Treg therapy to date have shown limited efficacy. To understand how Treg dosing impacts T1D progression, we formulate an extension of equations (25) - (31) to allow for the administration of Tregs into the lymph node compartment. We replace equation (27) with

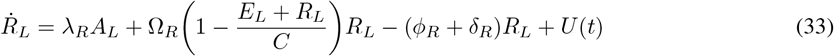

Where

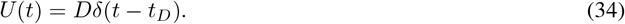

In (34), *t*_*D*_ denotes the time of dose, *D* is the number of dosed Tregs, and *δ* is the Dirac delta function, which models a bolus injection of regulatory T cells at time *t*_*D*_. All other equations remain the same, so that the system takes the form

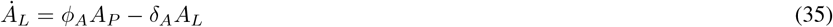

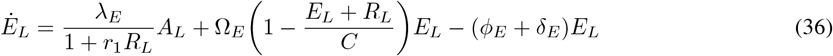

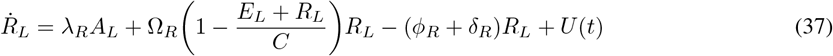

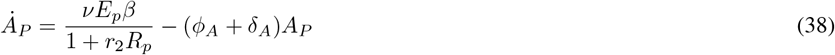

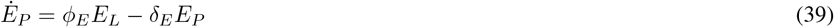

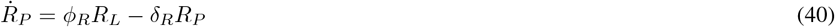

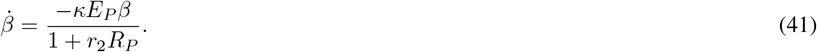

#### 4.1.4 Type 1 diabetes immune model with regulatory T cell dosing and APC depletion

While Treg therapy as a monotherapy has demonstrated limited clinical efficacy with respect to delaying the onset of T1D, Treg therapy in combination with antigen depletion therapy has yielded promising results. For example, a phase I/II trial in patients with new-onset T1D found that patients who received a combination of rituximab, which binds to CD20 on B cells and can elicit apoptosis, and Treg therapy responded better than those receiving either rituximab or Treg monotherapies [48]. To better understand how Treg dosing/APC depletion combination therapy works, we extend system (35)-(41) to incorporate APC depletion at the time of Treg dosing. For simplicity, we assume that APCs are fully depleted at the time of Treg dosing, so that when *t* ≥ *t*_*D*_ (see equation (34); *t*_*D*_ is the time at which Tregs are administered), equations (36) and (37) are replaced with

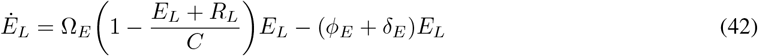

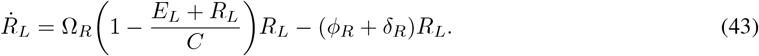

Outside of this extension, model equations (35) - (41) remain unchanged. Note that the only difference in the (*E*_*L*_, *R*_*L*_) equations (when compared to the original T1DI model (35) - (41)) is that the APC component of the vector field is removed; we are thus assuming an idealized therapy in which APCs are instantaneously suppressed at the time *t*_*D*_ of Treg administration.

### 4.2 Parameter estimation

In this section, we present a detailed discussion of the parameter values employed in the simulations of the T1DI model. We note that when estimating parameters, we report both human and mouse estimates with the assumption that mouse estimates yield a reasonable approximation to human counterparts; this is not uncommon when parameterizing mathematical models [50].

1. Parameter *ϕ*_*A*_ represents the rate at which activated APC cells migrate from tissue (pancreas) to the lymph nodes. We utilize rates associated with dendritic cells (DCs) as they are among the most common APCs [55]. Once a DC is matured, it takes approximately 18 hours for it to appear in the lymph nodes (18 hours = 0.75 days) [50]. Thus the rate at which DCs travel from the pancreas to the lymph nodes is fixed as *ϕ*_*A*_ = ln(2)*/*0.75 ≈ 0.9242 day^−1^.
2. Parameters *ϕ*_*E*_ and *ϕ*_*R*_ represent the respective rates at which effector T cells and Tregs travel from the lymph nodes to the pancreas. Using estimates obtained in [50], we set *ϕ*_*E*_ = *ϕ*_*R*_ = 1 × 10^−3^ day^−1^.
3. Parameters *δ*_*A*_, *δ*_*R*_, *δ*_*E*_, and *δ*_*P*_ represent the natural decay rates of APC, Tregs, effector T cells, and peptide, respectively. The lifespan of effector T cells and Tregs has been estimated to be approximately 4-5 days, which we assume are similar for APCs [19]. Thus *δ*_*A*_ = *δ*_*R*_ = *δ*_*E*_ = 0.25 day^−1^. Peptide decays more rapidly, with *δ*_*P*_ = 1 day^−1^ [25].
4. Parameters Ω_*E*_ and Ω_*R*_ represent the proliferation rates of effector T cells and Tregs, respectively. T cells have been estimated to proliferate 2-3 times per day [19]. Thus we obtain the following lower and upper bounds for Ω_*E*_ and Ω_*R*_: ln(2)*/*(1*/*2) ≈ 1.39 day^−1^ and ln(2)*/*(1*/*3) ≈ 2.08 day^−1^. For simulations, we fix these rates as the mean of these estimates to obtain Ω_*E*_ = Ω_*R*_ = 1.735 day^−1^.
5. Parameters *λ*_*E*_ and *λ*_*R*_ represent activation rates of effector and regulatory T cells, respectively. We assume that activation occurs via a pool of resting cells through binding with an APC in the lymph nodes. Thus, when estimating *λ*_*E*_ and *λ*_*R*_, we must consider both the number of resting T cells as well as the contact rate of a regulatory or effector T cell with an APC. Following [19], we fix the resting pool for both Tregs and effector T cells to be 5 cells, and the contact rate with APCs as 0.2 day^−1^ for both regulatory and effector T cells. This therefore yields *λ*_*E*_ = *λ*_*R*_ = 1 day^−1^.
6. Parameter *C* represents the immunological carrying capacity of the lymph nodes. Utilizing the spleen as an approximate for all lymph nodes, we note that a typical mouse spleen contains 1 × 10^8^ splenocytes, approximately 80% of which are T cells [56]. Hence, we fix the carrying capacity as *C* = 0.8 × 1 × 10^8^ = 2.5 × 10^7^ cells.
7. Parameter *α* represents the accumulation rate of peptide. This rate has previously been estimated as *α* = 1 × 10^−5^day^−1^ [25].
8. Parameter *κ* represents the killing rate of beta cells by effector T cells. This rate has previously been estimated as *κ* = 0.14 × 10^−6^ day^−1^ [25].
9. Parameter *σ* represents the activation of APC by peptide. The authors of [19] estimate the activation of APC per antigen molecule (denoted therein as *G*) to be 0.0025 × 10^−4^. We note that the units for peptide considered in this work are “cells”, and we thus assume a scaling factor *s >* 0 such that *G* = *sP*; this corresponds to each peptide “cell” being comprised of a large number of peptide molecules. We arbitrarily fix this scaling factor *s* to be 1 × 10^6^ − 1 × 10^10^, as the initial condition for *G* used in [19] is on the order of 10^8^. Thus, we obtain the following lower and upper bounds for *σ* =: 0.0025 × 10^−4^ × 10^6^ = 0.25 and 0.0025 × 10^−4^ × 10^10^ = 2500.
10. Parameters *r*_1_ and *r*_2_ represent scaling factors for the effect of Treg mechanisms on both activation of effector T cells by APC and the killing of beta cells by effector T cells, respectively. The values for both *r*_1_ and *r*_2_ are arbitrarily chosen to be between 0 − 0.1.

### 4.3 Regulatory T cell quality

We provide a higher resolution discretization of Figure 2a in Figure 9. Note that the transition between disease onset time *t*_*c*_ for small versus large *ν* is continuous, but that a rapid transition occurs for *ν* ≈ 1 × 10^−4^.

**Figure 9.**
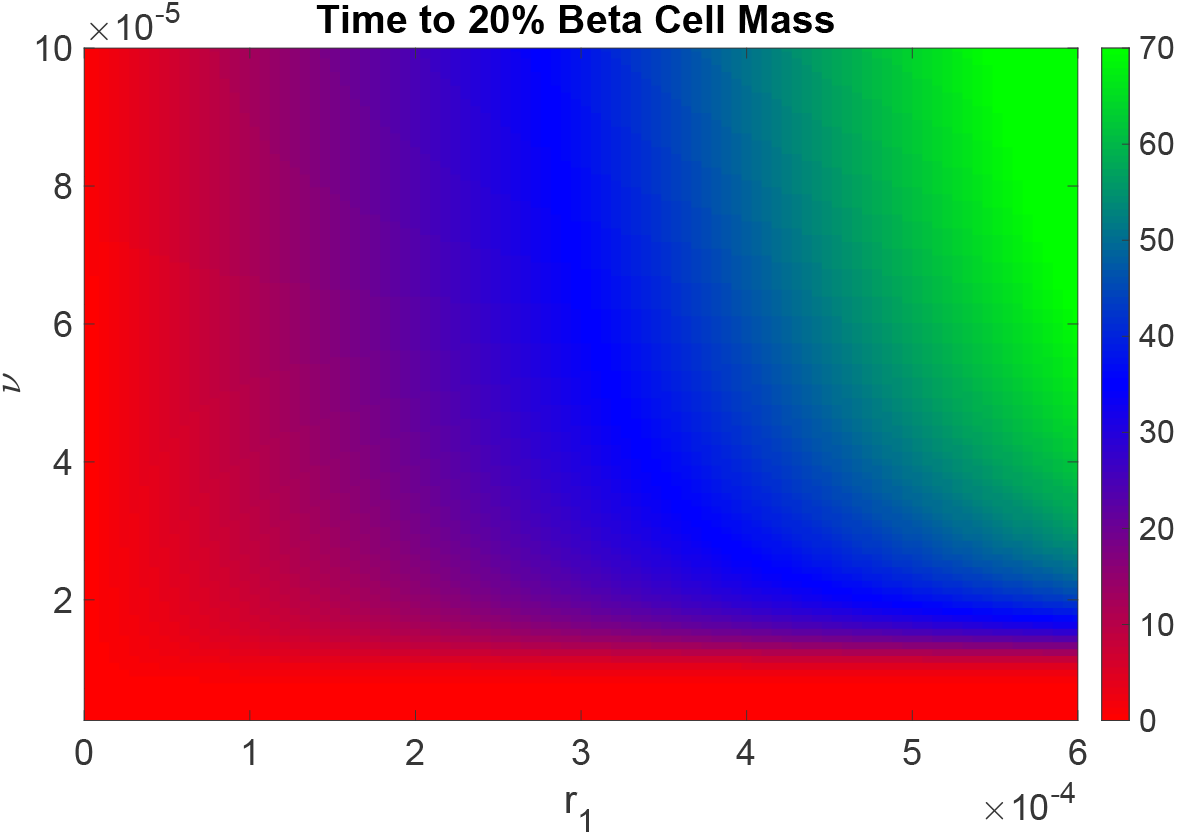
Time *t*_*c*_ to 20% beta cell mass of the T1DI model when varying *ν* and *r*_1_ while setting *r*_2_ = 0. Initial conditions utilized are *R*_*L*_(0) = *E*_*L*_(0) = 10 and *B*(0) = 1 × 10^6^, while other initial populations are 0. All other parameters may be found in Table 2. Treg therapy is not applied, so that *U* (*t*) ≡ 0. Red indicates early disease onset, blue indicates late disease onset, and green indicates no disease onset. Compare to Figure 2a.

**Figure 10.**
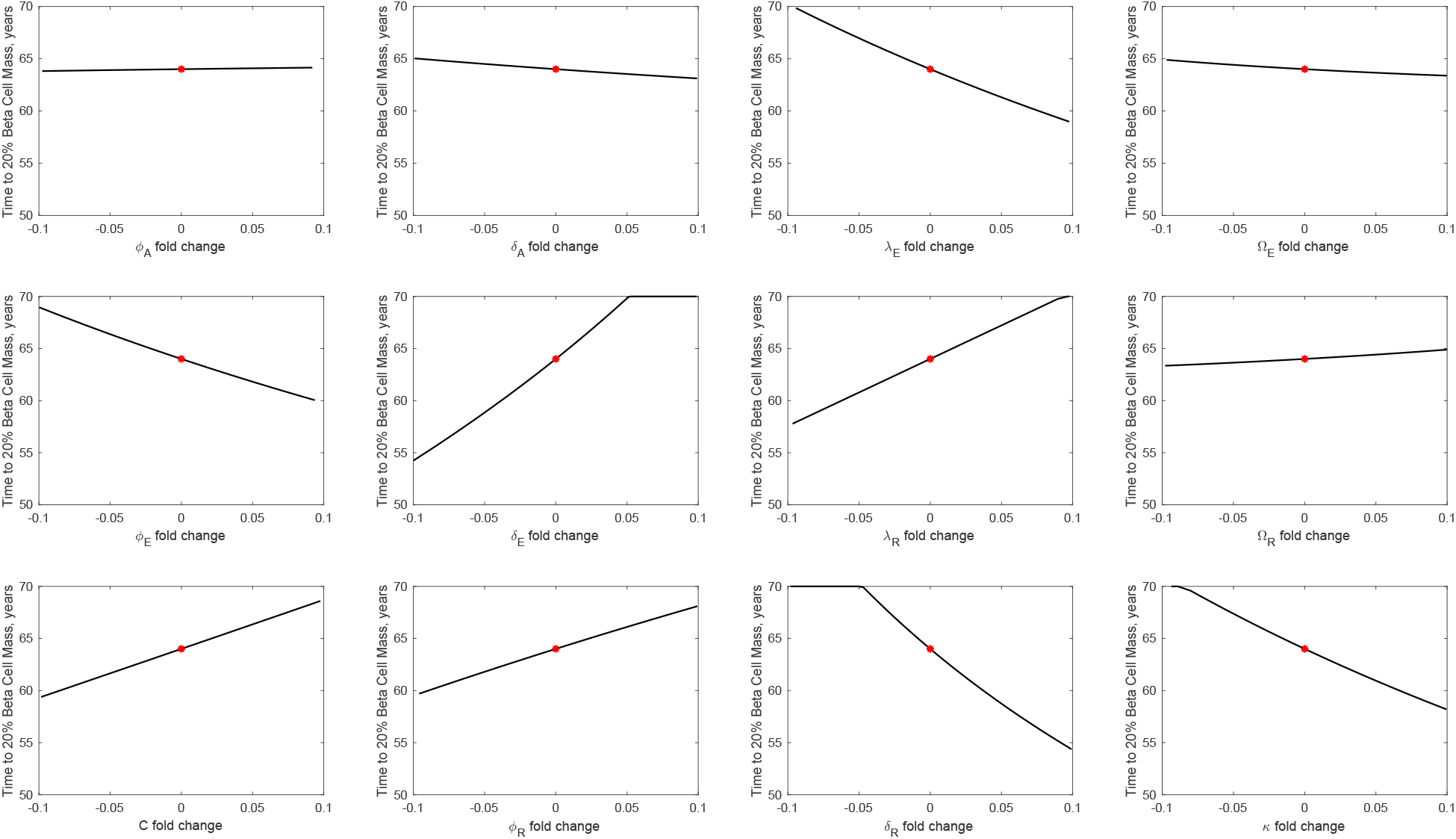
Parameter sensitivity analysis for the T1DI model. Here, *ν* = 0.01 and *r*_1_ and *r*_2_ correspond to late disease onset parameters (*r*_1_ = 0.8 × 10^−5^ and *r*_2_ = 1 × 10^−5^). Initial conditions utilized are *R*_*L*_(0) = *E*_*L*_(0) = 10 and *B*(0) = 1 × 10^6^, while other initial populations are 0. All other parameters may be found in Table 2. We compute *t*_*c*_ (see equation (8)) as a function of the varied parameter, with all other parameters fixed as in Table 2. The lower and upper bounds for each parameter is taken to be ± 10% its value reported in Table 2. We take 100 random samples between the upper and lower bound for each parameter and calculate the time *t*_*c*_ to 20% beta cell mass.

**Figure 11.**
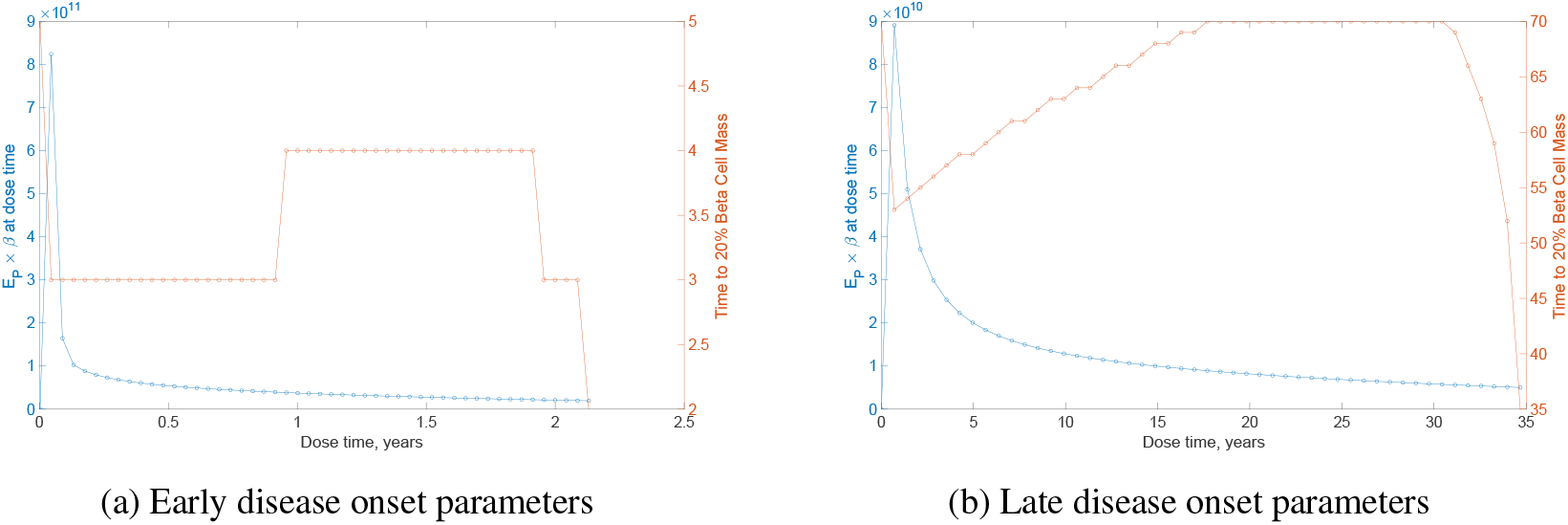
Plot of dose time (x-axis, *t*_*D*_), *E*_*P*_ × *β* (left coordinate, y-axis, blue), and time to 20% beta cell mass (right coordinate, y-axis, orange) for *ν* = 1 × 10^−5^ for (a) early disease parameter values (*r*_1_ = 0.2 × 10^−5^ and *r*_2_ = 0.2 × 10^−5^) and (b) late disease parameter values (*r*_1_ = 0.8 × 10^−5^ and *r*_2_ = 1 × 10^−5^). For all simulations, the dose is fixed at *D* = 1 × 10^9^ cells. Initial conditions are *R*_*L*_ = *E*_*L*_ = 10 and *B* = 1 × 10^6^, while other initial populations are 0. All other parameters may be found in Table 2.

**Figure 12.**
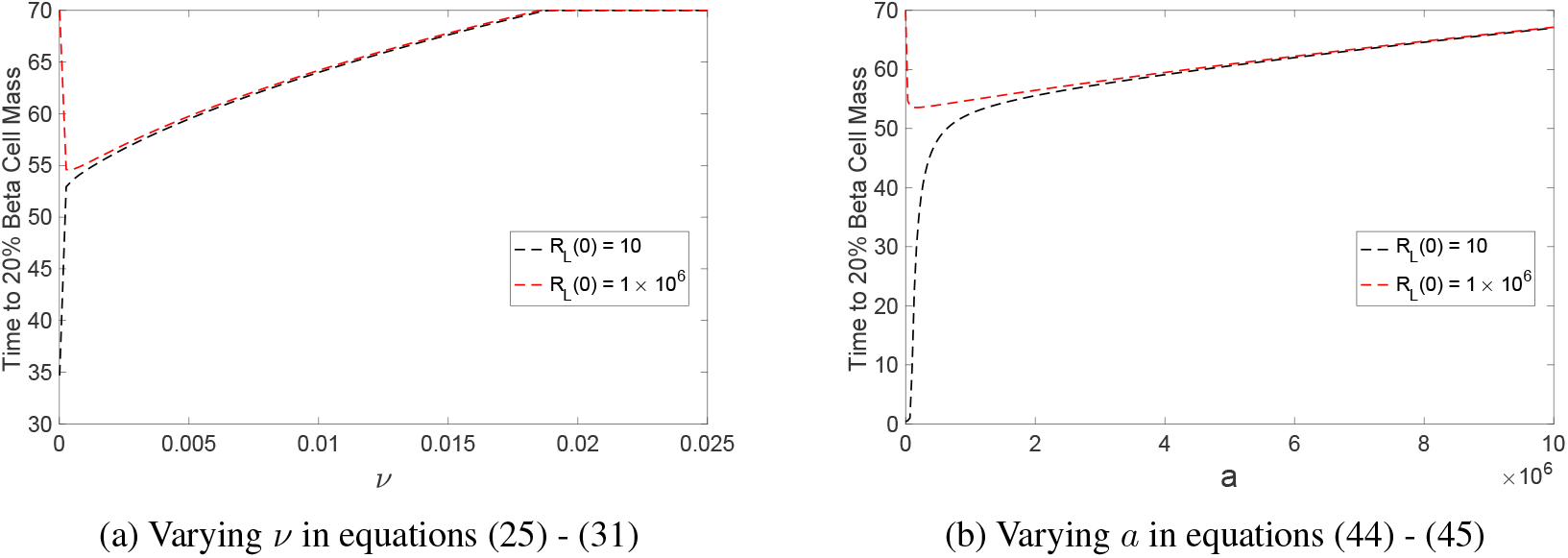
Time to 20% beta cell mass for *R*_*L*_(0) = 10 (black) and *R*_*L*_(0) = 1 × 10^6^ when (a) varying *ν* in equations (25) - (31), and (b) varying *a* in equations (44) - (45). Here, *E*_*L*_(0) = 10 and *β*(0) = 1 × 10^6^, and we fix late disease onset parameters (*r*_1_ = 0.8 × 10^−5^ and *r*_2_ = 1 × 10^−5^). All other initial populations are 0, and the remaining parameters may be found in Table 2.

### 4.4 Sensitivity analysis

We perform a sensitivity analysis on all parameters fixed throughout simulations (i.e. all parameters except *r*_1_, *r*_2_, and *ν*). Sensitivity is computed with respect to time *t*_*c*_ to 20% initial beta cell mass, and is measured as the slope of *t*_*c*_ with respect to fold changes in the corresponding parameter. Results are provided in Figure 10. We determined that *δ*_*E*_ and *δ*_*R*_ were among the most sensitive parameters, with *t*_*c*_ increasing rapidly with respect to *δ*_*E*_, while time to disease decreases with respect to *δ*_*R*_. Furthermore, we also observe that the system is sensitive to *λ*_*E*_ and *λ*_*R*_: *t*_*c*_ decreases with respect to *λ*_*E*_, and increases with respect to *λ*_*R*_. These parameters further suggest that the relative fitness of regulatory versus effector T cells is important to the dynamics of the T1DI model, as the sensitive parameters are all related to (*E*_*L*_, *R*_*L*_) interactions.

### 4.5 Optimal dosing time

We discuss the optimal Treg dosing times observed for *ν* = 1 × 10^−5^, as provided in Figures 4 and 5 in Section 2.3; note that we are not considering APC depletion in this section. Figure 11 provides the dose time *t*_*D*_ (x-axis) versus both *E*_*P*_ × *β* at *t*_*D*_ (left coordinate, y-axis) and time *t*_*c*_ to 20% beta cell mass (right coordinate, y-axis). In Figure 11a we observe that for early dose times, *E*_*P*_ × *β* at *t*_*D*_ is low, and thus dosing is effective at delaying T1D onset (time to disease is 5 years as opposed to the no Treg dosing time of ≈ 2 years). However, as *E*_*P*_ × *β* at dose time grows and peaks near 0.2 years, dosing is much less effective. Following the peak in *E*_*P*_ × *β* at *t*_*D*_, we note that disease onset *t*_*c*_ increases as *E*_*P*_ × *β* at dose time decreases. Finally, we observe that after approximately 2 years, dosing Tregs is too late to be effective to significantly decrease *t*_*c*_, despite lower *E*_*P*_ × *β* at *t*_*D*_. In Figure 11b we see similar results, with *t*_*D*_ between 15 to 30 years achieving a no disease state.

### 4.6 Impact of APC dynamics

We explore APC’s role as related to the impact of *ν* on disease progression and response to regulatory T cell therapy. We consider a simplified T1DI model, where we replace equations (26) and (27) with

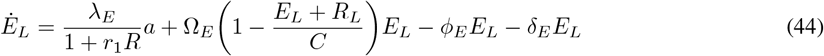

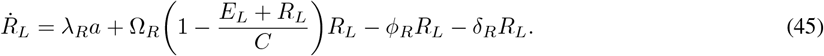

Here *a* represents a *constant* APC population, where we do not distinguish between APC in the lymph nodes (*A*_*L*_) and pancreas (*A*_*P*_), so that the APC population is fixed globally as the constant parameter *a* ≥ 0. We thus ignore APC dynamics (equations (25) and (28)), and consider the modified system (44) - (45) together with (28) - (31). Our goal is thus to understand the connection between *ν* and APC levels, specifically with regard to responses to Treg therapy; recall that *ν* regulates APC via (28). Figure 12 demonstrates that varying a constant APC population *a* in equations (44) - (45) yields a qualitatively similar dynamic with respect to Treg therapy to varying *ν* in equations (25) - (31); note that for simplicity, Treg therapy is modeled as an increase in *R*_*L*_(0), as opposed to the more general *U* (*t*) considered in Section 4.1.3. Precisely, we observe that individuals with a small *ν* parameter exhibit a significant response to Treg therapy, while those with large *ν* values yield essentially no improvement in time to disease for increased *R*_*L*_(0) values (Figure 12a). A completely analogous response occurs in the constant APC model, as observed in Figure 12b. That is, when varying the constant APC population *a*, we see that small *a* values respond to Treg therapy, while large *a* values do not. Hence we conclude that varying the parameter *ν* effectively varies the amount of APC in the system, which suggests that APC dynamics are essential when designing Treg therapies. This further motivates the investigation of combining APC depletion and Treg administration to increase time to disease *t*_*c*_.

This phenomenon can be understand theoretically via competition between effector and regulatory T cells with constant external activation. We consider the following model:

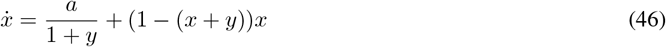

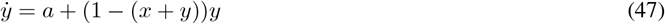

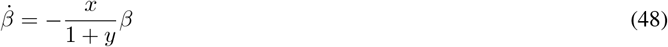

Here we ignore biophysical compartments (lymph nodes and pancreas) as well as APC dynamics; as in (44) - (45), APC is assumed fixed at a constant non-negative population *a* ≥ 0. In relation to the T1DI model, state *x* corresponds to the effector T cell population, *y* the regulatory *T* population, and *β* the beta cell mass. Note that although the above is technically a three-dimensional system, the *x* and *y* dynamics are uncoupled from *β*, so that the latter can be considered as simply an output of the two-dimensional (46) - (47) system, which can be analyzed via standard phase plane techniques. Our goal is to understand the role of the value of *a* in time to disease for different initial Treg populations. As in the original model (25) - (31), time to disease onset is quantified when *β* cell mass decreases beyond a critical threshold (see equation (8)). Lastly, we note that we have normalized all other parameters to be one for simplicity, as they do not affect the qualitative dynamics which wish to demonstrate with this simplified model.

We consider the two-dimensional system (46) - (47) in the first quadrant (*x, y* ≥ 0), as *x* and *y* correspond to (simplified) cell populations; note that the first quadrant is forward invariant. For any fixed *a >* 0, we can solve for the *x* and *y* nullclines as a function of *x*:

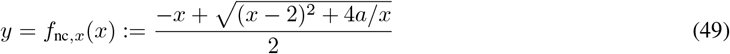

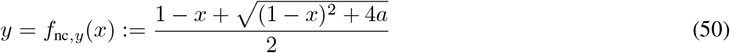

We make the following observations:

1. *f*_nc,*x*_ and *f*_nc,*y*_ are both decreasing as a function of *x*.

2. The nullclines satisfy the following limits:

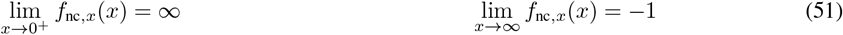

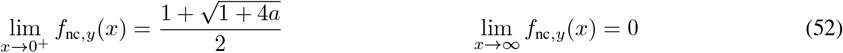

3. There exists a unique steady state 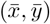 of system (46) - (47).

The existence/uniqueness of the steady state 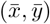 can be deduced by noticing that the *x* and *y* nullclines intersect uniquely in the first quadrant. The coordinates of the steady state satisfy

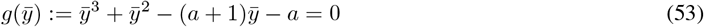

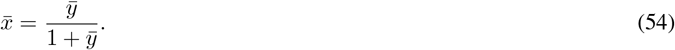

Since *g*(0) =− *a <* 0, *g*^*′*^(0) = − (*a* + 1) *<* 0, and the discriminant of *g*^*′*^ is positive, *g*^*′*^(*y*) = 0 has exactly one solution, so that *g* has one and only one positive root 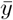. Thus for 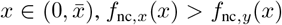 while the reverse is true for 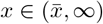. By analyzing the sign of 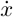 and 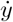 along the *y* and *x* nullclines respectively, we are able to deduce the directions of motion of trajectories in the first quadrant, and conclude that 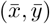 is globally asymptotically stable. A representative plot of nullclines, the steady state, directions of motion, and two sample trajectories (small and large *y*(0), which model Treg therapy) are provided in Figure 13.

**Figure 13.**
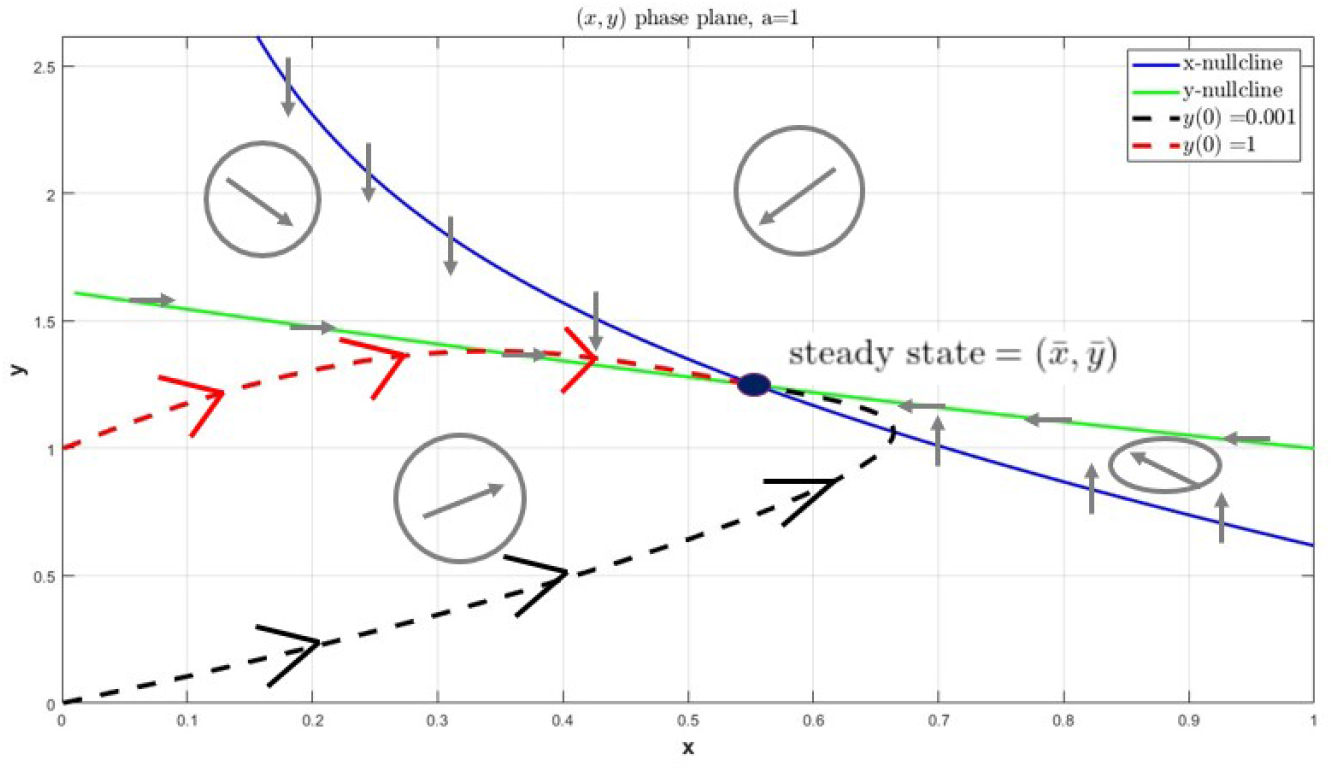
Nullclines (blue and green), trajectories (black and red), steady state (purple), and directions of motion (grey) for system (46) - (47). Here *a* is fixed as 1, but the same essential geometry/dynamics exist for all *a >* 0. Note that all solutions asymptotically converge to the steady state (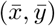), and that the different *y* initial conditions mimic regulatory T cell therapy as in Figure 12. In both trajectories, the *x* initial conditions are fixed at *x*(0) = 0.001.

We now investigate the change in response, i.e. time *t*_*c*_ to critical beta cell mass, as a function of *a*. We simulate system (46) - (48) for three different values of *a*: *a* = 0.01 (small), *a* = 1 (intermediate), and *a* = 100 (large). Results are provided in Figure 14, with rows corresponding to different constant *a* values, and columns providing the (*x, y*) phase plane (left), beta cell dynamics (middle), and (*x, y*) time series dynamics (right). As in the full T1DI model, we measure outcomes by time to disease *t*_*c*_, which is the time at which beta cell mass drops reaches 20% of its initial value (*β*_*c*_ in the middle column of Figure 14). For small values of *a* (Figure 14a), we observe a large increase in time *t*_*c*_ to disease onset by increasing the number of regulatory T cells (*y*): *t*_*c*_ ≈ 9 when *y*(0) = 0.001, while *t*_*c*_ ≈ 38 when *y*(0) = 1. As *a* increases, this difference in *t*_*c*_ decreases. Specifically, we observe in Figure 14b that *t*_*c*_ ≈ 6 for *y*(0) = 0.001, while *t*_*c*_ ≈ 8 for *y*(0) = 1 for *a* = 1, and when *a* = 100 in Figure 14c, *t*_*c*_ ≈ 20 for both initial conditions. This behavior is completely analogous to what observed in Figure 12, where we observe a large difference in time to disease onset for different initial Treg populations when *ν* (Figure 12a) or *a* (Figure 12b) is small; this difference then decreases to 0 as the respective parameter is increased. Hence the efficacy of Treg therapy can be understood as a function of APC levels.

**Figure 14.**
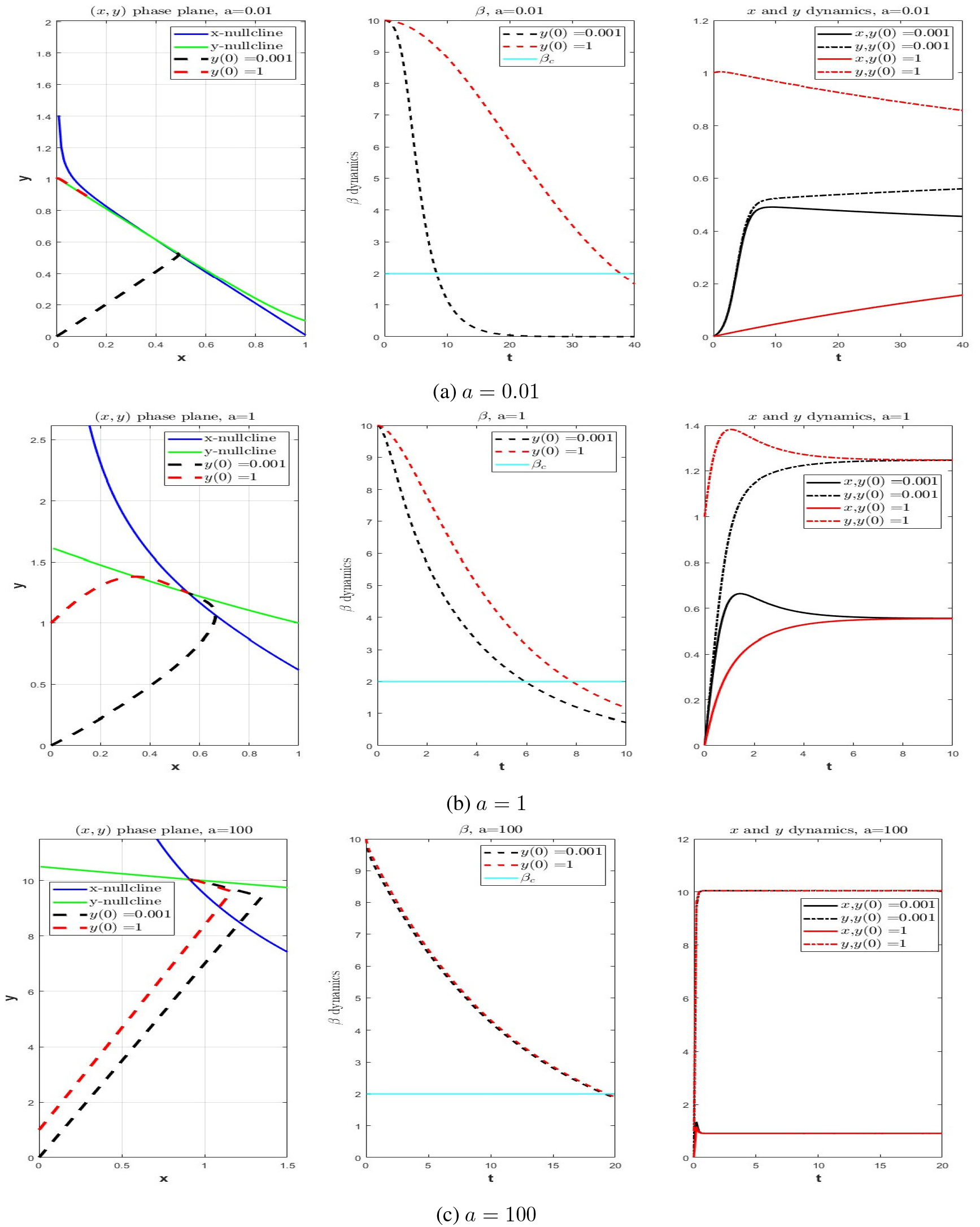
Simulation of simplified immune model (46) - (48) for different constant APC values *a*. The (*x, y*) phase plane dynamics are provided in the left column subplots, with the middle and right columns containing *β* and *x, y* solution curves, respectively. Initial conditions for *y* vary (black and red), and initial conditions for *x* are fixed at *x*(0) = 0.001 in all simulations.

We note that the dynamics observed in Figure 14 can be understood via standard perturbation theory arguments [57]. When *a* is small, system (46) - (47) takes the form of a regular perturbation problem (*ϵ* = *a*), whose solution can be approximated (up to order one) by the solution of the unperturbed system

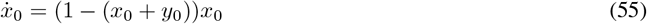

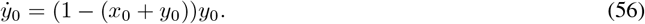

Solutions of system (55) - (56) remain on the lines 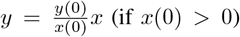 determined by the initial conditions (*x*(0), *y*(0)), and the system possesses an infinite number of fixed points defined by *x* + *y* = 1; note that each fixed point is neutrally stable, and not attracting. For *y*(0) = 1, *x*(0) = 0.001, the solution trajectory is near this line, and hence the vector field is close to 0 ∈ ℝ^2^, so that the trajectory moves more slowly compared to trajectories with initial conditions away from this line (e.g. *y*(0) = *x*(0) = 0.001). Analytically, we see from system (46) - (47) that the *x* and *y* nullclines coalesce to form this line of fixed points as *a* → 0^+^, so that for small positive *a*, the nullclines nearly coincide, which thus creates a region in phase space where the vector field is small (see the left panel of Figure 14a). The initial conditions with large *y*(0) (red) are near this region, and hence the corresponding solution moves slowly to the (global) steady state 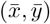, so that significantly longer time is required for beta cell mass to decay, in comparison with the small *y*(0) solution. This can be seen easily from the expression for 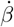 in (48), as larger *y* and smaller *x* inhibit beta cell decay. See the right panel in Figure 14a for a comparison of trajectory rates of convergence to steady state for *y*(0) small (black) and large (red), as well as the middle panel for the corresponding beta cell response.

As *a* increases, the nullclines of system (46) - (47) separate, causing the *y*(0) large (red) trajectory to increase in speed, so that *β*_*c*_ is reached more quickly (*t*_*c*_ ≈ 38 in Figure 14a, while *t*_*c*_ ≈ 8 in Figure 14b). Large *a* dynamics can be understood again using a perturbation approach. We rewrite the equation in standard form, by defining

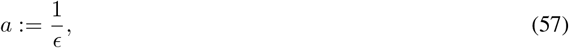

where 0 *< ϵ* ≪ 1. Multiplying by *ϵ*, system(46) - (47) takes the form

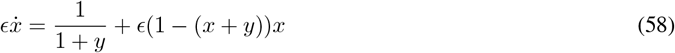

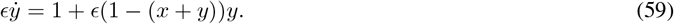

The form of the above system suggests that there exists an inherent time scale *τ* of the form

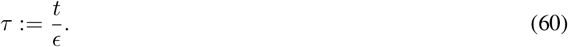

With respect to this *τ* time scale, i.e. defining states

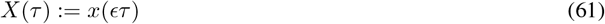

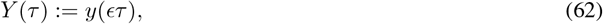

we observe that dynamics of *X* and *Y* are given by the following ODE system:

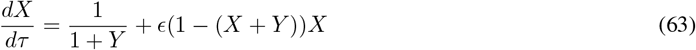

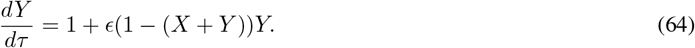

Note that by the change of variables, we have transformed the original singular perturbation problem (*x* and *y*) into a regular perturbation problem (*X* and *Y*). For small *ϵ*, the dynamics of the (*X, Y*) system can be approximated (up to order one) by the system

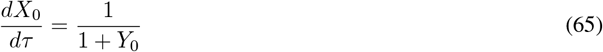

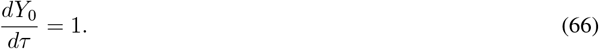

The dynamics of (65) - (66) are clear: each component increases for all initial conditions, with the rate of increase of *X*_0_ inhibited by the increasing *Y*_0_ values. Of course, such an increase cannot can not be sustained indefinitely in the full system (63) - (64), as the carrying capacity eventually forces convergence to the global steady state 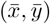 as discussed previously. However, the time scale for this convergence is with respect to *τ*, which by (60) is on the the order of 1*/ϵ*. Hence, convergence to this unique steady state is fast for small *ϵ*, which implies all trajectories, regardless of initial conditions, rapidly approach steady state, and thus exhibit near-identical beta cell dynamics. Thus, when *a* is large (*ϵ* is small), there is essentially no difference in time to disease *t*_*c*_ between small and large *y* initial conditions, and thus there is little effect of Treg therapy. This is demonstrated both in the original model in Figure 12, as well as in the simpler system (46) - (48) in Figure 14c, where the trajectories in the latter are observed to be nearly identical after a short initial transient (see middle and right panel specifically in Figure 14c).

### 4.7 Stability analysis

We provide the basic dynamical properties of model (25) - (31). For convenience, we provide the ODE system below:

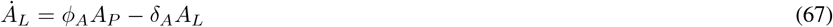

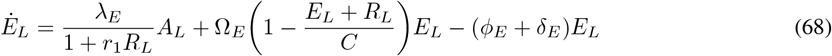

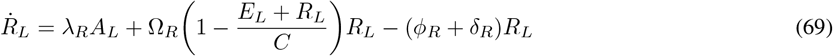

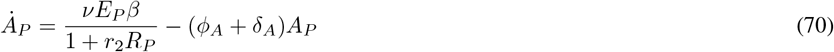

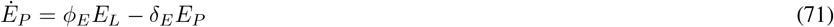

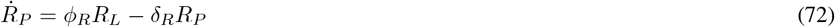

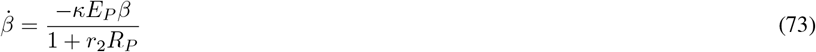

We begin with a technical lemma, which will be utilized in the proofs of Theorems 1 and 2. The following says that under relatively mild assumptions, a decaying time-varying perturbation does not affect the convergence of a system with globally attracting steady-state.

#### Lemma 1.

*Consider the autonomous ODE system*

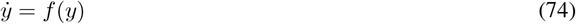

*together with its perturbation by a time-varying function g:*

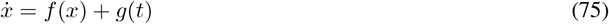

*Here f is assumed to be locally Lipschitz vector field on an open connected subset U of* ℝ^*n*^, *g* : ℝ→ ℝ^*n*^ *is continuous, and the non-perturbed system* (74) *has a globally attractive steady state L. Suppose that*

*1*. 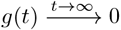, *and*

*2. the solutions of the perturbed system* (75) *are bounded*.

*Then the solution of any IVP associated to* (75) *also converges to L as t* → ∞.

*Proof*. By assumption, *L* is an equilibrium of *f* (*f* (*L*) = 0). By the change of coordinates *y* → *y* − *L*, we may assume that *L* = 0 is a globally attractive steady state of *f*, and that *U* is a open set containing 0. By Lyapunov’s stability theorem, it will be sufficient to construct a Lyapunov function for the perturbed system (75) [58]. That is, it will be enough to show that there exists a positive definite 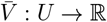 such that for *t* sufficiently large,

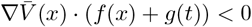

for all *x* ∈ *U \*{0}. Since 0 is globally asymptotically stable for the non-perturbed system (74), converse Lyapunov theory implies the existence of a smooth Lyapunov function *V* for *f* [59]. Thus,

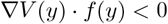

for all *y* ∈ *U \* {0}. We claim that *V* is the desired 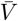. Note that

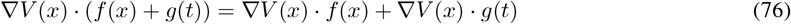

By the Cauchy-Schwarz inequality,

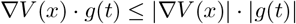

Since solutions of (75) are assumed bounded and *V* is smooth, |∇*V* (*x*)| is bounded along trajectories. As *g*(*t*) → 0 as *t* → ∞, we have that

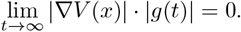

Note that this statement is true along trajectories of *f* (*x*) + *g*(*t*), as solutions *x* were assumed bounded. Thus for *t* sufficiently large, ∇*V* (*x*) · *f* (*x*) *<* 0 for all *x* ∈ *U \* {0} implies that

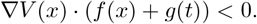

Hence, *V* is a Lyapunov function for the perturbed system (75), and thus all solutions converge to *L* = 0, as desired.□

Our next results states that all solutions of (67) - (73) remain non-negative and bounded for all times *t* ≥ 0. Denoting the state of the system in ℝ^7^ as

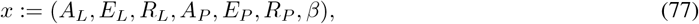

this implies that the unique solution to the corresponding initial value problem (IVP)

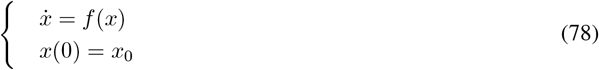

exists for all times *t* ≥ 0 when 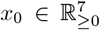 we are interested in nonnegative initial conditions since all variables represent cell counts. Here the vector field 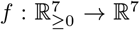 is defined via the ODE system (67) - (73). Note that *f* is smooth, and hence standard existence-uniqueness results apply.

#### Proposition 1.

*Solutions x*(*t*) *of the IVP* (78) *with non-negative initials conditions remain bounded and non-negative for all t* ≥ 0.

*Proof*. Denote the solution at time *t* of the IVP associated to (78) with initial condition *x*_0_ as *x*(*t, x*_0_) ∈ ℝ^7^; each component is similarly denoted *x*_*i*_(*t*), and denote each component of *f* as *f*_*i*_. By examining the form of *f*, we see that if *x*_*i*_(*t*) = 0, then *f*_*i*_(*x*(*t*)) ≥ 0. Thus, the set 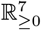 is invariant with respect to the flow of the vector field *f*.

To show that solutions remain bounded, we first note that *β* is non-increasing and bounded below by 0, and thus converges. Hence *β* is bounded (by *β*(0)). Define

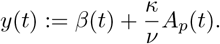

Using (70) and (73), we see that

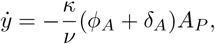

which implies that *y* is converges, and hence is bounded. As *β* is bounded, this further implies that *A*_*P*_ is bounded. Equation (67) then implies that *A*_*L*_ is bounded. Indeed, if *M* is a bound for *A*_*P*_, then we see that *A*_*L*_ is bounded above by 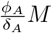 as 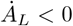 if 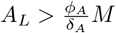. By an analogous argument, *E*_*L*_ and *R*_*L*_ are bounded, as are *E*_*P*_ and *R*_*P*_, completing the proof.□

#### Theorem 1.

*For positive parameter values and initial conditions we have that the antigen-presenting cells A*_*L*_ *and A*_*P*_ *converge to* 0. *Furthermore, the beta cell population β converges to a value between β*(0) *and* 0.

*Proof*. To prove the above statements, we will utilize Barbalat’s Lemma [60]. For convenience, we provide the result here. Assume that a differentiable function *h* : ℝ→ ℝsatisfies the following:

i. 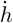 is uniformly continuous, and
ii. *h* has a limit at infinity.

Then we can immediately conclude that

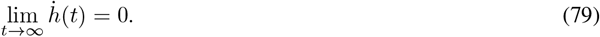

As solutions of ODEs are absolutely continuous [43], *A*_*L*_, *A*_*P*_, and *β* are absolutely, and hence uniformly, continuous. Hence to apply Barbalat’s Lemma, we will only need to verify condition (ii) of the above.

We have previously shown (see the proof of Proposition 1) that *β* converges, as it is a bounded below and decreasing function.

We now prove that *A*_*P*_ converges to 0. By Barbalat’s Lemma, since *β* converges, we know that 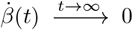. Equation (73) then implies that

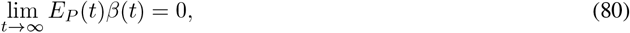

as *E*_*P*_ is bounded by Proposition 1. We now apply Lemma 1, with

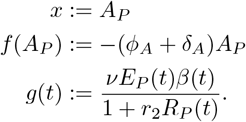

As 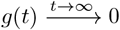 and solutions of 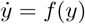 converge to 0, Lemma 1 implies that *A*_P_ also converges to 0. An analogous argument implies that *A*_*L*_ converges to 0. Specifically, let *x* = *A*_*L*_, *f* (*A*_*L*_) = −*δ*_*A*_*A*_*L*_, and *g*(*t*) = *ϕ*_*A*_*A*_*P*_ (*t*), and again apply Lemma 1 to conclude that 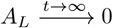.

We now state and prove our main theorem concerning the asymptotic behavior of (67) - (73). In words, it says that for all initial conditions, the system converges to a steady state, and that unless a delicate balance in growth/death rates exists between effector and regulatory *T* cells (*k*_*E*_ = *k*_*R*_ below), there is no coexistence between different types of immune cells.

#### Theorem 2.

*Consider the system* (67) *-* (73), *and define*

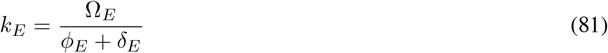

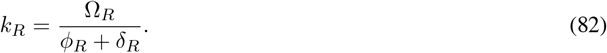

*Assume that k*_*E*_, *k*_*R*_ *>* 1; *note that a value smaller than* 1 *would imply monotonic decay of the respective type of immune cell, which is not biologically relevant for type 1 diabetes. Let x*(*t*) ∈ ℝ^7^ *denote the state of the system at time t* ≥ 0 *(as in* (77)*). For positive parameter values and initial conditions, the asymptotic behavior of x may be classified as follows:*

*1. If k*_*R*_ *> k*_*E*_,

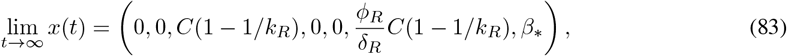

*where β*_∗_ ∈ [0, *β*(0)). *That is, A*_*L*_, *E*_*L*_, *A*_*P*_, *and E*_*P*_ *are all asymptotically vanishing, and the steady state value of beta cells β*_∗_ *is determined by the initial conditions of the system*.

*2. If k*_*R*_ *< k*_*E*_,

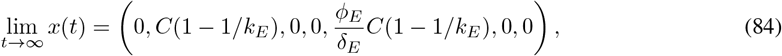

*That is, A*_*L*_, *R*_*L*_, *A*_*P*_, *R*_*P*_, *and β are all asymptotically vanishing. Note that in this case, β cells are asymptotically eliminated*.

*3. If k*_*E*_ = *k*_*R*_,

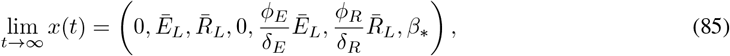

*where Ē*_*L*_ *and* 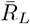 *lie on the line* 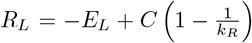, *and β*_∗_ ∈ [0, *β*(0)); *Ē*_*L*_ *and* 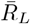 *are dependent on the initial conditions of the corresponding IVP. Furthermore, if Ē*_*L*_ *>* 0, *then β*_∗_ = 0.

*Proof*. We have already shown that *A*_*L*_ and *A*_*P*_ approach 0 in Theorem 1, and furthermore that *β* converges to *β*_∗_. Assume first that *k*_*R*_ *> k*_*E*_. We again apply Lemma 1, with

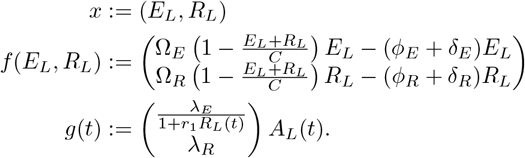

A standard phase plane analysis (e.g. analyzing nullclines) shows that *k*_*R*_ *> k*_*E*_ implies that solutions of 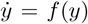 satisfy

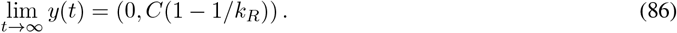

Since 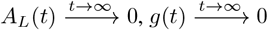 and thus Lemma 1 together with boundedness (Proposition 1) implies that

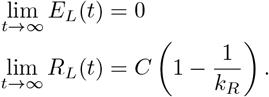

Since *E*_*L*_ converges to 0, another application of Lemma 1 implies that

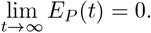

To see the limiting value for *R*_*P*_, note that we can integrate (72) to obtain

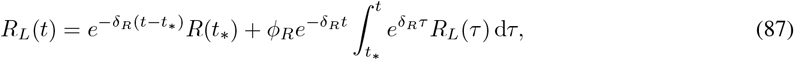

for any *t, t*_∗_ ≥ 0. Since 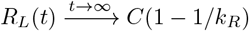 for any *ϵ >* 0, we can find *t*_∗_ *>* 0 such that

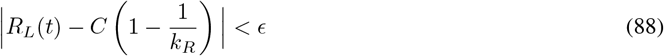

for all *t* ≥ *t*_∗_. Bounding the integral in the right-hand side of (87) via (88), we obtain the following upper bound for *R*_*L*_(*t*) on [*t*_∗_, ∞):

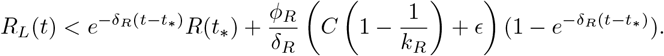

Thus we have the follow estimate on the limit superior of *R*_*L*_:

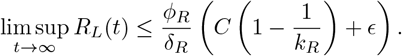

Since this holds for all *ϵ >* 0,

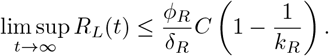

Applying the same technique to estimate the limit inferior of *R*_*L*_, we obtain

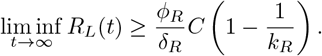

Hence

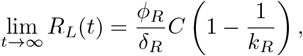

as claimed.

The proof for the case when *k < k* is analogous. Note that in this case, *β* = 0, since 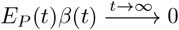 (see (80)), and in this case, *E*_*P*_↛0 as *t* → ∞

When *k*_*R*_ = *k*_*E*_, consider

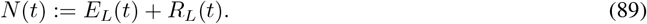

For notational simplicity, denote

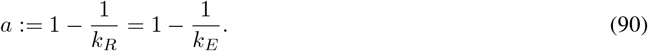

Using (68) and (69), we see that *N* satisfies

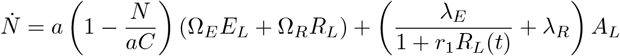

We have already seen that 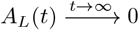, and hence we can again apply Lemma 1, with

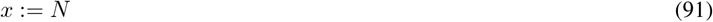

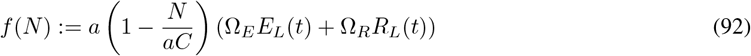

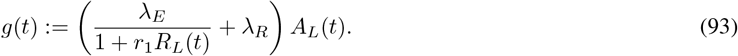

We note that we technically need an extension of Lemma 1 to nonautonomous vector fields *f* (*t, x*), since *f* in (92) is nonautonmous; however such an extension is immediate by the theory of nonautonomous Lyapunov functions. Since *E*_*L*_(*t*), *R*_*L*_(*t*) *>* 0 for all *t* ≥ 0 (positive invariance of for trajectories with positive steady states), we see that the solution 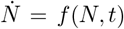 is monotone and bounded, and thus converges. Hence Lemma 1 implies that the solution of 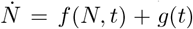 also converges, i.e. that *E*_*L*_(*t*) + *R*_*L*_(*t*) converges to a point on the line 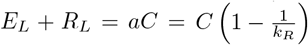 as desired.

